# Genetic population structure and demographic history of Pacific cod in Japanese waters: Implications for stock identification using SNP markers

**DOI:** 10.64898/2026.03.11.710969

**Authors:** Akira S. Hirao, Kay Sakuma, Tetsuya Akita, Satoru N. Chiba

**Affiliations:** Fisheries Resources Institute, Japan Fisheries Research and Education Agency, 2-12-4 Fukuura, Kanazawa-ku, Yokohama, Kanagawa 236-8648, JAPAN; Fisheries Resources Institute, Japan Fisheries Research and Education Agency, 1-5939-22 Suido-cho, Chuo-ku, Niigata, Niigata 951-8121, JAPAN; Fisheries Resources Institute, Japan Fisheries Research and Education Agency, 2-4-1 Nakanoshima, Toyohira-ku, Sapporo, Hokkaido 062-0922, JAPAN

**Keywords:** Genetic stock identification, GRAS-Di, Population genomics, PSMC, Whole-genome sequencing

## Abstract

Pacific cod is a key species in North Pacific fisheries, and its stock assessment and management units are separated according to biological, geographical, and administrative information. Understanding the fine-scale genetic population structure of this species is crucial for effective management, particularly in regions such as Japan, where complex coastal geography and localised fisheries management prevail. Therefore, in this study, we analysed genome-wide single nucleotide polymorphisms (SNPs; 6,035 loci) in 496 individuals of Pacific cod sampled from 33 sites around the Japanese archipelago via genotyping by random amplicon sequencing-direct (GRAS-Di) analysis. Our analyses revealed three major genetic groups: Japanese Broad Range, Northernmost Honshu–Hokkaido (NHH), and Western Sea of Japan groups. These groups exhibited significant genetic differentiation (global *F*_ST_ = 0.056), distinct levels of nucleotide diversity, and group-specific genome-wide patterns of Tajima’s D. Moreover, demographic history reconstruction based on whole-genome sequencing of three representative individuals revealed that each genetic group followed distinct demographic trajectories since the last glacial period. Importantly, the NHH group, related to the Mutsu Bay spawning aggregation and previously shown to exhibit strong natal homing in tagging surveys, was genetically identified for the first time in this study. Isolation-by-distance was observed across Japanese waters and within the Japanese Broad Range group, but not within the NHH group, suggesting that gene flow is generally restricted by geographic distance, except within the NHH group. To evaluate the potential for genetic stock identification, we extended a resampling-based cross-validation framework by incorporating outlier detection to assess marker selection strategies. Over 500 background SNPs were required to achieve >90% assignment accuracy for genetic stock identification, whereas only eight or more outlier SNPs showed comparable performance. These findings suggest that carefully selected SNP panels, particularly those including outlier loci, substantially improve stock discrimination. Overall, our study demonstrates the fine-scale genetic structure and demographic history of Pacific cod in Japanese waters and highlights the utility of practical marker strategies for enhancing the biological realism of fisheries assessment and supporting sustainable fisheries management.

## Introduction

Fish stocks, groups of individuals with similar demographic and genetic profiles, are managed based on the best available scientific evidence, while considering environmental, economic, and social sustainability. Ideally, stock identification should be grounded in detailed knowledge of the biological characteristics of the populations concerned. However, in practice, stock delineation is often based on geographical, socio-economical, or administratively defined boundaries because of the frequent lack of comprehensive biological data. The term “stock” is sometimes defined as “a marine biological resource that occurs in a given management area” (European Commission 2014), and stock identification remains a critical prerequisite for fishery assessment and management (Cadrin *et al*. 2023). Even when biological data are available, stock identification can be complicated by dynamic spatiotemporal processes, such as the mixing of populations with distinct spawning groups during feeding migrations (e.g., Berg *et al*. 2017, Seljestad *et al*. 2024). To address these challenges, population genomics analysis has emerged as a powerful tool for delineating stocks and resolving stock-specific contributions to mixed-stock fisheries in marine ecosystems (Bernatchez *et al*. 2017, Benestan 2019).

Pacific cod (*Gadus macrocephalus*, Tilesius 1810), widely distributed along the continental shelf of the North Pacific Ocean, is among the most important species in North Pacific fisheries (Ketchen 1961, Bakkala 1984). It is also among the most extensively studied marine fishes in terms of population genetics (Saitoh 1998, Canino *et al*. 2010, Kim *et al*. 2010, Suda *et al*. 2017, Drinan *et al*. 2018, Sakuma *et al*. 2019, Smirnova *et al*. 2019, Spies *et al*. 2020, Spies *et al*. 2021, Fisher *et al*. 2022, Spies *et al*. 2022b). For Pacific cod fisheries, stock assessment and management units are separated at various spatial scales based on biological, geographical, and administrative information. The largest-scale stock management is implemented in Alaska by the National Oceanic and Atmospheric Administration Fisheries and North Pacific Fishery Management Council. These agencies recognize three management stocks within the Alaska Exclusive Economic Zone (>2.3 million km²): the Gulf of Alaska, Aleutian Islands, and eastern Bering Sea (Barbeaux *et al*. 2022, Hulson *et al*. 2022, Spies *et al*. 2022a). Genome-wide single nucleotide polymorphism (SNP) analyses have largely supported these management units (Drinan *et al*. 2018, Spies *et al*. 2020), while also suggesting a more complex population structure with seasonal feeding migrations (Spies *et al*. 2020).

Pacific cod populations in the western Pacific are managed at finer spatial scales, particularly along the Japanese coasts, where complex island-chain geography at the continental margin has led to the delineation of management units across seven sea areas: Sea of Okhotsk, Sea of Japan Coast of Hokkaido, Nemuro Strait, North Pacific Coast of Hokkaido, North Pacific Coast of Honshu, Northern Sea of Japan Coast of Honshu, and Western Sea of Japan (WSJ) Coast of Honshu (Fujiwara *et al*. 2025, Hamabe *et al*. 2025, Ito *et al*. 2025, Kuwahara *et al*. 2025, Sakai *et al*. 2025, Sakuma *et al*. 2025) (Fig. 1; Table 1). The last unit has been delineated but not yet subjected to stock assessment. Previous genetic studies have reported that Pacific cod species in the WSJ are genetically distinct from those in other regions based on microsatellite and mitochondrial DNA analyses (Suda *et al*. 2017, Sakuma *et al*. 2019). However, local genetic groups of Pacific cod are expected not only in the WSJ but also along other Japanese coasts, where natal homing to spawning grounds, such as those in Mutsu Bay in northern Honshu (Fukuda *et al*. 1985, Miura *et al*. 2019), may contribute to population differentiation. Therefore, applying high-resolution genomic approaches based on genome-wide SNP markers may reveal previously undetected population structures.

**Figure 1.**
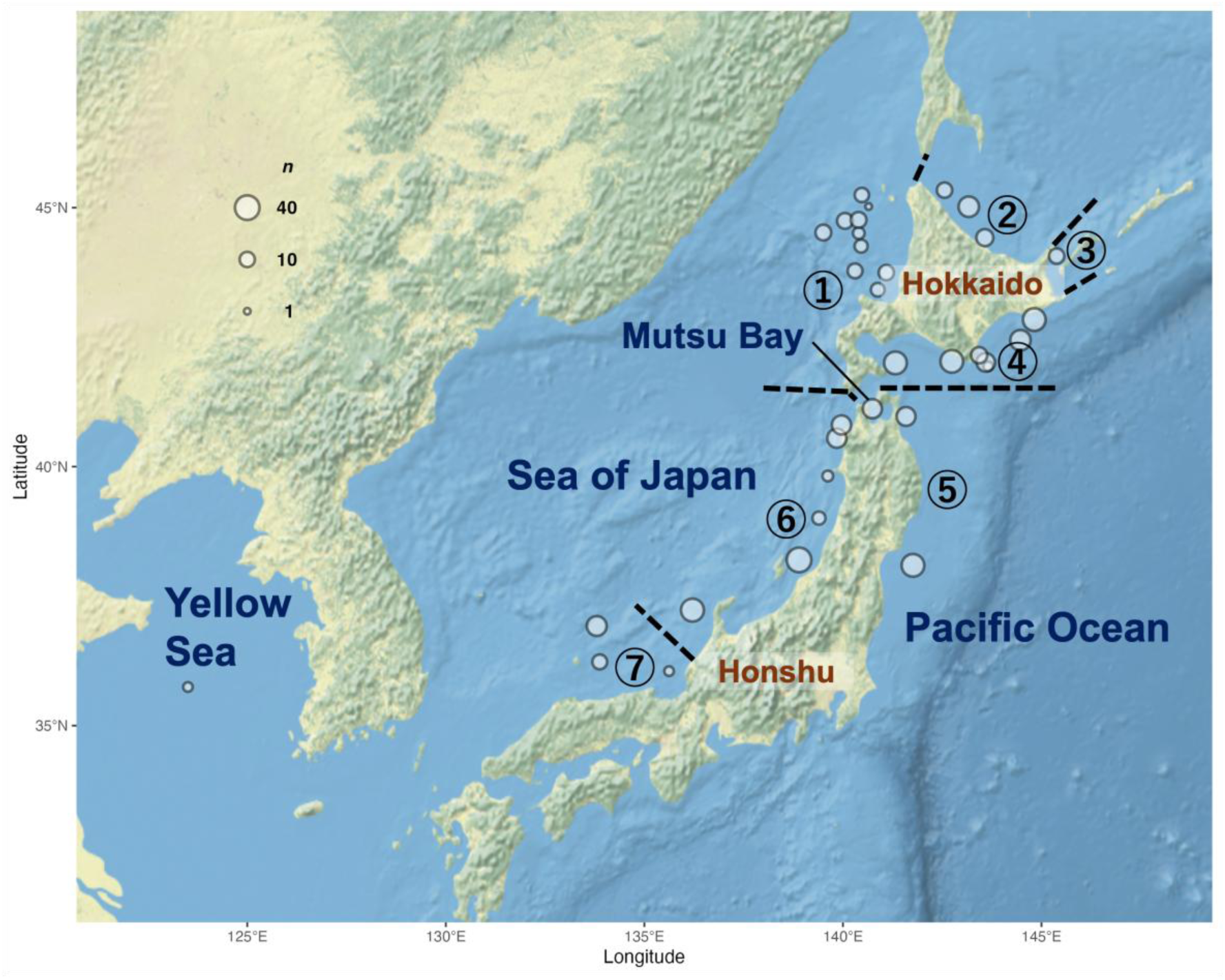
Sampling locations of Pacific cod genotyped in Japanese waters. Circle size indicates the sample size at each site listed in Table S1. Current management units are labelled with numbers corresponding to Table 1, and their boundaries are shown as dashed lines.

**Table 1.** Summary of sampling metadata for the genotyping by random amplicon sequencing-direct (GRAS-Di) dataset (see Table S1 for details and Fig. 1 for geographic context).

| Management unit | *N* site | N Pacific cod |  |
| --- | --- | --- | --- |
|  |  | Sequenced | Genotyped |
| 1. Sea of Japan Coast of Hokkaido | 10 | 78 | 78 |
| 2. Sea of Okhotsk | 3 | 48 | 48 |
| 3. Nemuro Strait | 1 | 10 | 10 |
| 4. North Pacific Coast of Hokkaido | 8 | 172 | 167 |
| 5. North Pacific Coast of Honshu | 2 | 51 | 50 |
| 6. Northern Sea of Japan Coast of Honshu | 6 | 122 | 110 |
| 7. Western Sea of Japan Coast of Honshu | 3 | 34 | 33 |

Interpreting the patterns of genetic population structure requires acknowledging that such structures are often dynamic rather than strictly static or homogenous. Population composition can vary across space and time, and commercial catches or survey samples may include individuals from multiple populations (e.g., Seljestad *et al*. 2024). Genetic stock identification (GSI), which assigns individual fish to source populations using genetic data, provides an effective means to detect immigration and quantify mixed-stock composition. Because genetic differentiation among marine fish populations is often subtle, GSI based solely on neutral SNPs typically requires several hundred markers, making it costly even when optimised SNP panels are used. Outlier markers potentially under selection have been incorporated to reduce the number of markers while maintaining discriminative power. However, commonly used outlier detection methods (e.g., *F*_ST_-based outlier tests) are prone to false positives. To address this issue, we extended a resampling-based cross-validation framework to jointly evaluate population structure and detect outlier SNPs in each iteration. Developing effective strategies for genetic marker selection to discriminate genetic groups can provide valuable insights into dynamic population structures in marine ecosystems.

This study focused on Pacific cod along the Japanese coasts, with three main objectives: (1) To elucidate the genetic population structure using genome-wide SNPs, (2) to reconstruct the historical demography of each identified genetic group using whole-genome sequencing (WGS) data, thereby providing evidence of independent population units, and (3) to develop a marker selection strategy for discriminating genetic groups for future applications in GSI.

## Materials and Methods

### Sampling, genotyping by random amplicon sequencing-direct (GRAS-Di) analysis, and SNP genotyping

Sampling locations of Pacific cod are shown in Fig. 1, and sample attributes are listed in Table 1 and Supplementary Table S1. In total, 515 samples from 33 sites were subjected to GRAS-Di, a reduced-representation sequencing method, using a two-step polymerase chain reaction library with random primers (Enoki *et al*. 2018). Detailed procedures for GRAS-Di and subsequent SNP genotyping are provided in the Supplementary Note. High-quality bi-allelic SNPs were called following the Genome Analysis Toolkit best practice pipeline (DePristo *et al*. 2011) using the chromosome-level reference genome assembly for *G. macrocephalus* (National Center for Biotechnology Information accession: GCF_031168955.1).

Three SNP datasets were generated for downstream analyses: Dataset 1 comprising bi-allelic SNPs (including singletons) and non-variants, which was used for estimating nucleotide diversity and analysing linkage disequilibrium (LD) decay, Dataset 2 comprising bi-allelic SNPs with minor allele frequency (MAF) > 0.05 after LD pruning (*r*^2^ < 0.1), which was used for population structure analyses, and Dataset 3, which was an extended version of Dataset 2 including two additional whole-genome sequenced samples from the Yellow Sea (DRA/SRA/ERA: SRR21531029 and SRR17394978). The detailed procedure for generating Dataset 3, which involved integrating genotypes of the GRAS-Di dataset (Dataset 2) with those of the two samples from the Yellow Sea, is provided in the Supplementary Note.

### Genetic structure and diversity

To investigate the genetic population structure of Pacific cod, we performed individual-based principal component analysis (PCA) and discriminant analysis of principal components (DAPC) using the R package “Adegenet” (Jombart 2008). Following Thia (2023), the DAPC model was constructed using *k*−1 PC axes, with the optimal *k* determined by Cattell’s rule, identifying the “elbow” point in the scree plot where eigenvalues start to level off. The geographic distribution of individuals assigned to genetic groups identified by DAPC (see section “Genetic structure and diversity” in Results) was visualised using the R package “mapmixture” (Jenkins 2024). Phylogenetic relationships among individuals were inferred using the maximum-likelihood method implemented in IQ-TREE v2.2.2.7 (Minh *et al*. 2020), applying the GTR+ASC model with 1,000 ultrafast bootstrap replicates (detailed procedures are provided in the Supplementary Note). Global and pairwise *F*_ST_ values among the identified genetic groups were calculated using the method described by Weir and Cockerham (1984), with confidence intervals (CIs) estimated via 1,000 bootstrap replicates.

Nucleotide diversity (π) within each identified genetic group was calculated based on Dataset 1 using 10-kb sliding windows as implemented in pixy v2.0.0 (Korunes and Samuk 2021, Bailey *et al*. 2025). The 95% CIs of π were estimated using bootstrapping windows with 5,000 replicates. Differences in π between groups were tested using the same bootstrapping procedure, with Bonferroni correction applied for multiple comparisons. To assess whether genome-wide LD varied among genetic groups, LD decay was evaluated based on the coefficient of determination (*r*^2^’) between SNP loci, with *r*^2^ adjusted for finite sample size (Waples 2024), as a function of chromosomal distance. Genome-wide distributions of Tajima’s D (Tajima 1989; equation 38) were assessed for each genetic group to infer population demographics. An unbiased estimate of Tajima’s D under missing genotype data (Bailey *et al*. 2025) was calculated using the original Python package ThetaRecov (https://github.com/akihirao/ThetaRecov).

Next, isolation by distance (IBD) was tested to determine the effect of geographic distribution on genetic differentiation. Least-cost paths (Adriaensen *et al*. 2003) between sampling sites, avoiding landmasses and shallow water areas (<10 m depth), were calculated using the R package “marmap” (Pante and Simon-Bouhet 2013). IBD was evaluated for sites with more than 10 genotyped individuals using the Mantel test, correlating pairwise *F*_*ST*_⁄(1 − *F*_*ST*_) values with geographic distances along least-cost paths.

### Demographic history

To infer the demographic history of each identified genetic group, we performed pairwise sequentially Markovian coalescent (PSMC) analysis (Li and Durbin 2011) using WGS data from one representative individual per group. The assignment tests described below were conducted using all background SNP markers to validate the representativeness of the individuals within their respective groups. WGS and PSMC analysis procedures are described in detail in the Supplementary Note.

### SNP marker selection for discriminating genetic groups

Practical SNP marker strategies were evaluated separately for background (before outlier analysis) and outlier SNPs (Fig. 2a and b, respectively). For potential application in GSI, we focused on two genetic groups outside the WSJ (see Results), where stock assessment has not yet been conducted. Assignment accuracy was assessed using a resampling-based cross-validation framework originally proposed by Chen *et al*. (2017) for their R package assignPOP, with custom modifications for outlier detection implemented using our own R scripts (https://github.com/akihirao/Gma_GRASDi). Resampling-based cross-validation approaches generally ensure independent training and test datasets (Anderson 2010) and minimise bias arising from unequal sample sizes among source populations (Wang 2017). Instead of using the machine-learning classifiers provided in assignPOP, we used DAPC to improve computational efficiency and maintain interpretability. Additionally, we incorporated customised procedures to reduce false positives in outlier detection, thereby enhancing the robustness of marker selection. Details of these procedures are provided below. In total, 465 samples (excluding those from the WSJ) and 6,035 SNPs (described below) were analysed.

**Figure 2.**
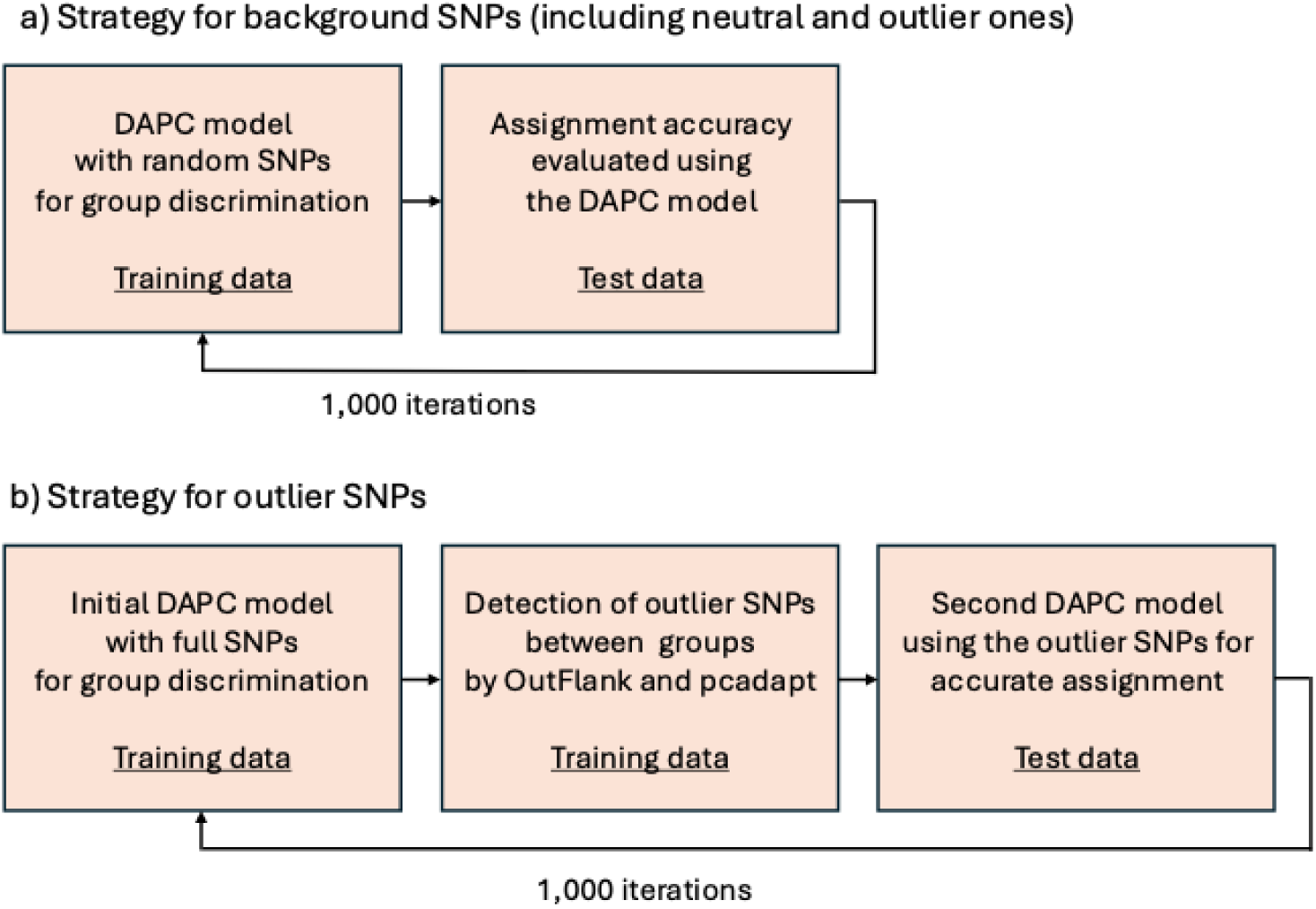
Workflow of resampling-based cross-validation for single nucleotide polymorphism (SNP) marker selection.

As illustrated in Fig. 2a, for background SNP markers (including both neutral and outlier markers), a resampling-based cross-validation with 1,000 iterations was conducted as described below. First, a DAPC model was constructed using resampled training data to maximise differentiation between the two genetic groups. In this procedure, 50 or 90% of the 465 samples were randomly assigned as training data, and the remainder were used as test data. Simultaneously, SNP markers were randomly selected at six levels (10, 100, 300, 500, 1,000, and all). Second, the trained DAPC model was applied to the test data using the corresponding SNP set. The estimated correct assignment rate (ECAR; Anderson 2010), defined as the proportion of correctly assigned individuals in the test data, was calculated using the assignment results from the full dataset as the reference.

For outlier SNP markers, resampling-based cross-validation with 1,000 iterations was conducted using a two-step DAPC modelling procedure with integrated outlier detection (Fig. 2b). First, an initial DAPC model was constructed using resampled training data to discriminate the genetic groups. In this procedure, 50 or 90% of the 465 samples were randomly assigned as training data, and the remainder were used as test data. All SNPs were retained in both datasets. Second, putative outlier SNPs were identified from the training data using two methods: OutFLANK v0.2 (Whitlock and Lotterhos 2015), a likelihood-based *F*_ST_ distribution model, and pcadapt v4.4.0 (Luu *et al*. 2017), a PCA-based method. In OutFLANK, locus-specific *F*_ST_ values between the two genetic groups identified in the previous step were calculated for all SNPs, and outliers were detected at a significance threshold of *q* = 0.05. In pcadapt, outliers were identified with an expected false discovery rate of 0.05 using an optimal *K* value of 2 (determined from the scree plot of the full dataset, with *K* values of 1–10). SNPs significantly detected by both OutFLANK and pcadapt were used in the next step. Third, a second DAPC model was constructed using only the outliers identified at six levels (top 3 based on *F*_ST_ estimates, top 5, top 8, top 10, top 20, and all) and was applied to the test data. ECAR was calculated using the assignment results from the full dataset as the reference. Finally, outlier SNPs consistently detected across cross-validation procedures with detection frequencies > 0.7 were functionally annotated using snpEff v5.1 (Cingolani *et al*. 2012) based on the reference genome annotation file (GCF_031168955.1-RS_2023_09).

## Results

### Identifying genome-wide SNP loci

GRAS-Di yielded 3.8 billion paired-end reads of 566.4 Gbp from 517 samples. After quality-filtering and alignment post-processing, an average of 4.7 M (±1.5 M SD) reads per individual were obtained. A variant-calling procedure with critical filtering excluded one sample because of its relatively low genotyping rate. Kinship analysis of the remaining 516 samples revealed one pair of genetically identical samples, indicating that the same fish had been mistakenly sequenced twice. Additionally, 19 samples exhibited unusually high multi-locus heterozygosity (*F* < -0.1) and elevated kinship, suggesting contamination during wet lab procedures. Consequently, one duplicate and 19 contaminated samples were excluded from further analysis. The remaining 496 GRAS-Di samples were genotyped at 2,321,838 loci, including 55,702 SNPs and 2,266,136 invariants (dataset 1), indicating a high genotyping rate of 90.3%. After further filtering, 6,035 SNPs were retained for population structure analyses by pruning for LD (*r*^2^ < 0.1) and an MAF threshold of 0.05 (Dataset 2). To assess the relationship with the waters surrounding the Korean Peninsula, the two Yellow Sea samples were added to Dataset 2, resulting in 6,030 SNPs (MAF > 0.05: LD-pruned; Dataset 3).

### Genetic structure and diversity

PCA of 496 individuals genotyped at 6,035 SNPs (Dataset 2) revealed three genetic groups along the PC1 and PC2 axes (Fig. 3; Supplementary Fig. S1). Based on Cattell’s rule applied to the PCA scree plot (Supplementary Fig. S2), DAPC was performed with the optimal number of clusters (*k* = 3), yielding posterior assignment probabilities exceeding 99.9% for each group (Fig. 4a). The geographic distribution of the following genetic groups is shown in Fig. 4b: Japanese Broad Range (JBR), Northernmost Honshu–Hokkaido (NHH), and WSJ groups. The JBR group was broadly distributed along Japanese waters, whereas most of the NHH group was regionally restricted to the Pacific side of Hokkaido and northernmost part of Honshu, including Mutsu Bay. The WSJ group occurred exclusively in the WSJ. PCA and DAPC analyses of Dataset 3, which included the two additional Yellow Sea samples, revealed the same three-cluster pattern as in Dataset 2 (Supplementary Figs. S4 and S5). The two Yellow Sea samples were assigned to the WSJ group in the DAPC model (Supplementary Fig. S5). A maximum-likelihood phylogenetic tree further supported the existence of the three genetic groups: JBR, NHH, and WSJ (Supplementary Fig. S6).

**Figure 3.**
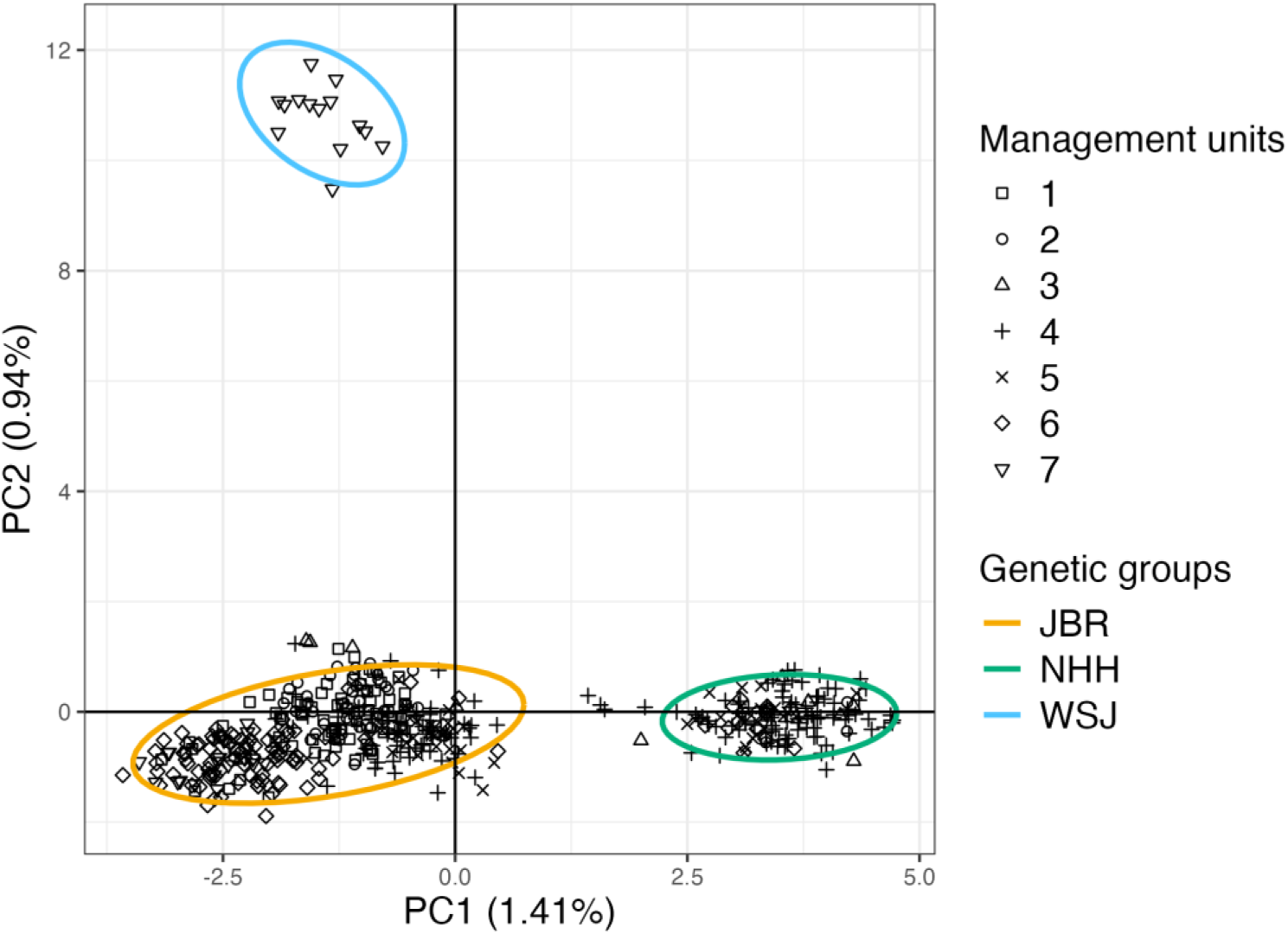
Principal component analysis (PCA) of 496 Pacific cod individuals sampled from Japanese waters. Different symbols correspond to management units, and coloured ellipses represent 95% confidence regions for the genetic groups inferred via discriminant analysis of principal components (DAPC).

**Figure 4.**
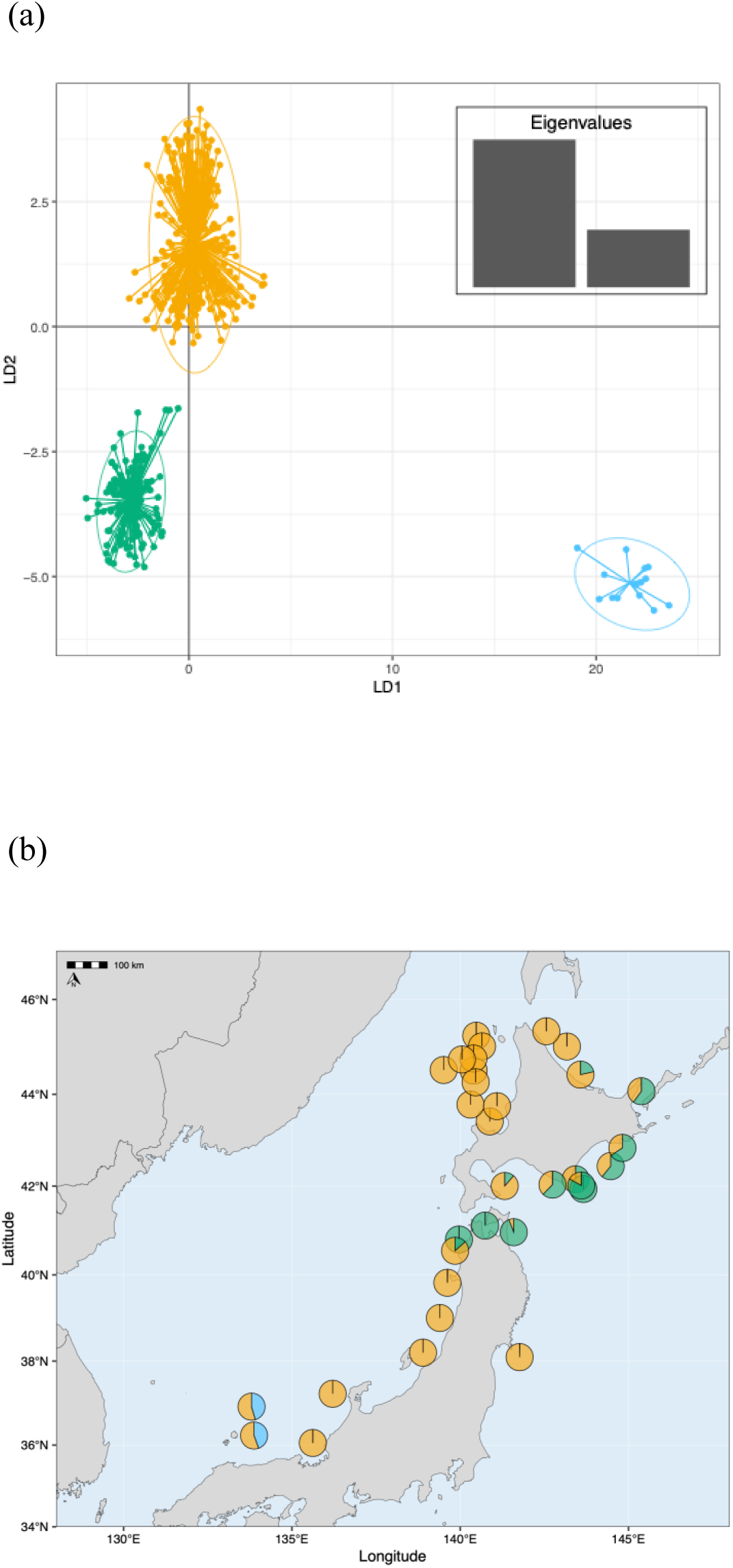
Genetic structure of Pacific cod in Japanese waters inferred via DAPC. (a) DAPC scatterplot showing three genetic groups. (b) Geographic distribution of groups represented by pie charts of membership proportions. The following colours indicate the groups identified via DAPC: Yellow = Japanese Broad Range (JBR) group, green = Northernmost Honshu–Hokkaido (NHH) group, and blue = Western Sea of Japan (WSJ) group.

Nucleotide diversity (π) values were 0.00118 (95% CI: 0.00115–0.00121), 0.00112 (0.00109–0.00115), 0.00107 (0.00104–0.00110), and 0.00117 (0.00114–0.00120) for JBR, NHH, WSJ, and overall, respectively (Table 2; Supplementary Fig. S7). The levels of π differed significantly between the JBR and WSJ groups (*P* < 0.05; window-blocked bootstrap test with Bonferroni correction); however, no other significant pairwise differences were detected. The global *F*_ST_ value was 0.056 (95% CI: 0.052–0.062), indicating significant differentiation among groups. Pairwise *F*_ST_ values varied: The lowest differentiation was observed between the JBR and NHH groups (*F*_ST_ = 0.0159), whereas high differentiation was noted between the JBR and WSJ groups (*F*_ST_ = 0.0608) and between the NHH and WSJ groups (*F*_ST_ = 0.0735). Moreover, the JBR and NHH groups exhibited low LD, with LD decaying below *r*^2^ = 0.03 within 8 kbp (Supplementary Fig. S8). In contrast, the WSJ group showed a rapid initial decay of LD below *r*^2^ = 0.1 within the first 10 kbp, but *r*^2^ remained above 0.055 across the entire range (up to 400 kbp), suggesting population subdivision within this group. Genome-wide Tajima’s D was negatively biased and differed significantly among groups (*P* < 0.001 for all pairs; using a linear mixed model with Student’s t errors [glmmTMB], accounting for windows-block structure with Holm adjustment; Table 2; Supplementary Fig. S9). The negative skew was most pronounced in the JBR group, followed by the NHH and WSJ groups. Genetic distance (*F*_ST_) plotted against geographic distance (Fig. 5) revealed significant IBD across Japanese waters and within the JBR group, but not within the NHH group (*r* = 0.44, 0.69, and 0.32, respectively, and *P* < 0.001, < 0.001, and = 0.08 for overall, JBR, and NHH, respectively). Notably, IBD was not tested for the WSJ group due to sampling of only two locations.

**Figure 5.**
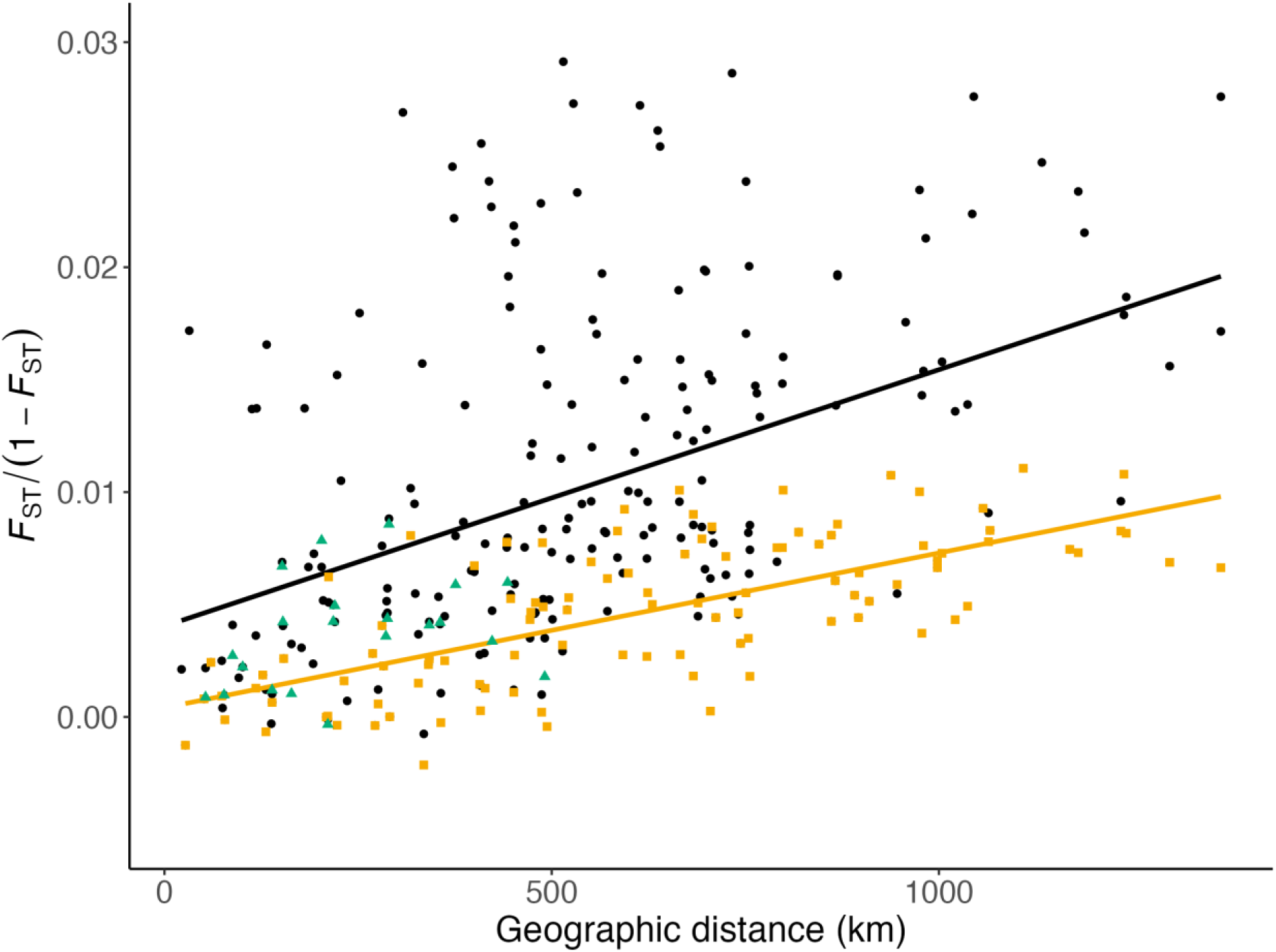
Isolation-by-distance pattern of Pacific cod in Japanese waters. A significant Mantel test correlation (*P* < 0.001) was observed for the overall samples (black) and JBR group (JBR: yellow), but not for the NHH group (green).

**Table 2.** Diversity statistics for the genetic groups of Pacific cod in Japanese waters.

| Genetic group | Nucleotide diversity (mean [95% confidence intervals]) | Tajima's D<br>(median [95% confidence intervals]) |
| --- | --- | --- |
| JBR | 0.00118 (0.00115–0.00121) | –1.645 (–1.679 to –1.617) |
| NHH | 0.00112 (0.00109–0.00115) | –0.956 (–1.005 to –0.920) |
| WJS | 0.00107 (0.00104–0.00110) | –0.477 (–0.546 to –0.407) |
| Overall | 0.00117 (0.00114–0.00120) | –1.667 (–1.699 to –1.633) |

### Demographic history of genetic groups

WGS reads were obtained from three individuals, one representing each of the three genetic groups. The average total base count per sample was 48.1 Gbp (38.3–61.0 Gbp), with an average sequencing depth of 50.0× (40.1–65.3×). Assignment tests confirmed that the three individuals were representative of their respective groups (*P* > 0.99).

The PSMC plot revealed that each genetic group experienced distinct demographic histories since the last glacial period (LGP, 116–11.7 ka according to Kukla *et al*. 2002; Fig. 6). The three groups shared a predation period during the LGP. After divergence during the LGP, they followed different demographic trajectories but showed a gradual population decline, followed by recent rapid expansion. The JBR group exhibited the smallest decline and most pronounced expansion, whereas the WSJ group showed the greatest shrinkage and least expansion, and NHH showed an intermediate pattern. Recent historical *N*_e_ ranked JBR > NHH > WSJ. Sensitivity analyses using generation times of 4 and 14 years, corresponding to the minimum and maximum values, with 9 years considered the most plausible estimate, confirmed that divergence occurred during the LGP (Fig. 6; Supplementary Fig.S10).

**Figure 6.**
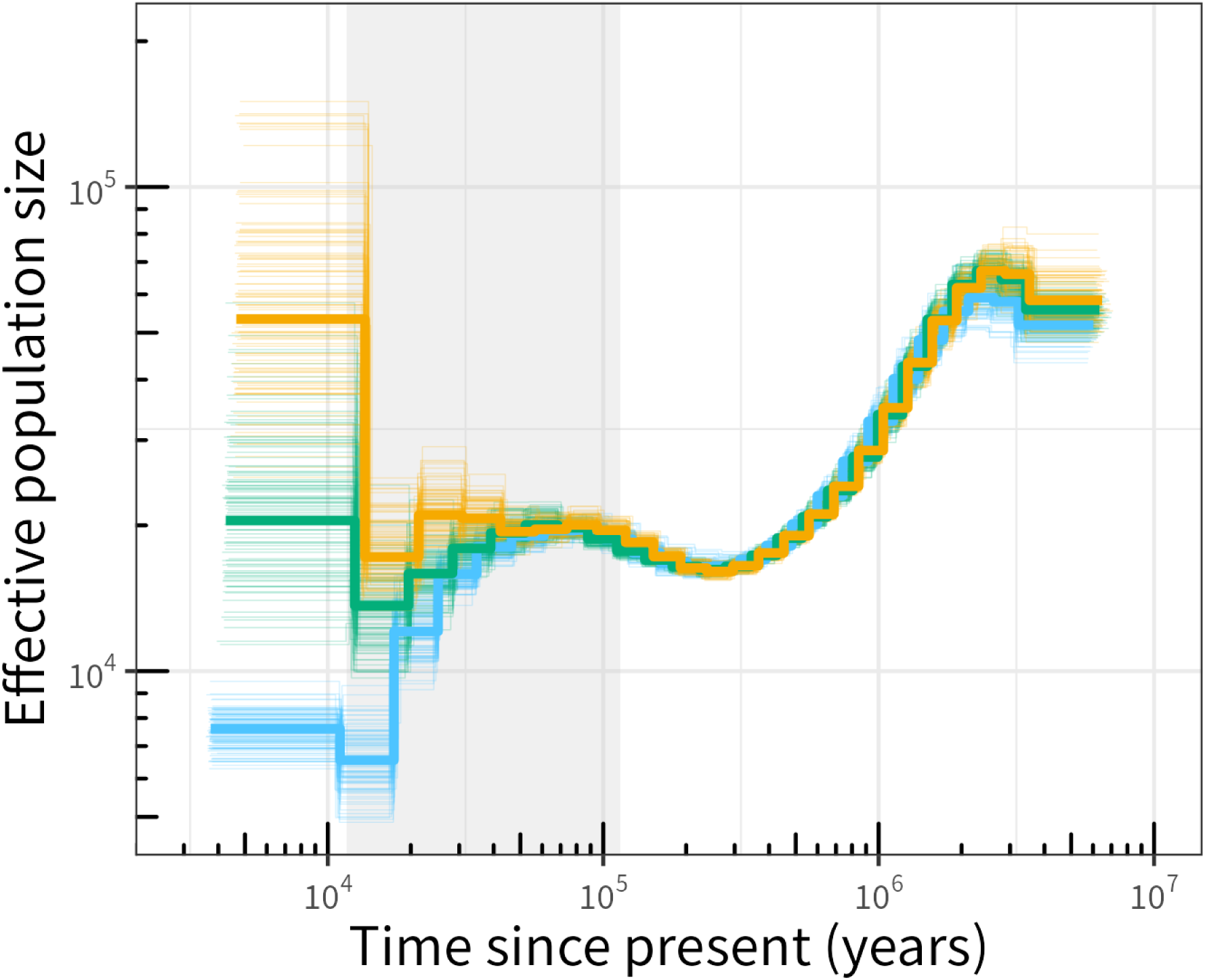
Pairwise sequentially Markovian coalescent (PSMC) estimates based on whole-genome sequencing (WGS) data from one representative individual of each genetic group: JBR (yellow), NHH (green), and WSJ (blue). Bold lines show estimates from observed data, and thin lines represent 100 bootstrap replicates. The shaded region corresponds to the last glacial period (LGP, 116–11.7 ka according to Kukla *et al*. 2002).

### Genetic assignment accuracy

Resampling cross-validation indicated that more than 500 background SNP markers were required to achieve GSI between the JBR and NHH genetic groups, with an ECAR exceeding 90% (Supplementary Fig. S11). In contrast, cross-validation incorporating outlier detection achieved over 80% ECAR using the top three outlier markers and over 90% ECAR using eight or more markers (Fig. 7). Among the 151 outlier SNPs detected across all cross-validation procedures, 8 and 18 SNPs were consistently identified from 50 and 90% of the training data, respectively, with detection frequencies > 0.7 (i.e., each SNP was identified as an outlier in at least 70% of the 1,000 resampling trials; Supplementary Fig. S12; Table S2). Of the 18 consistently identified outliers, 12 SNPs were located within introns or near protein-coding genes, three were located upstream or within introns of non-coding RNA genes, and the remaining three were located in intergenic regions (Supplementary Table S2).

**Figure 7.**
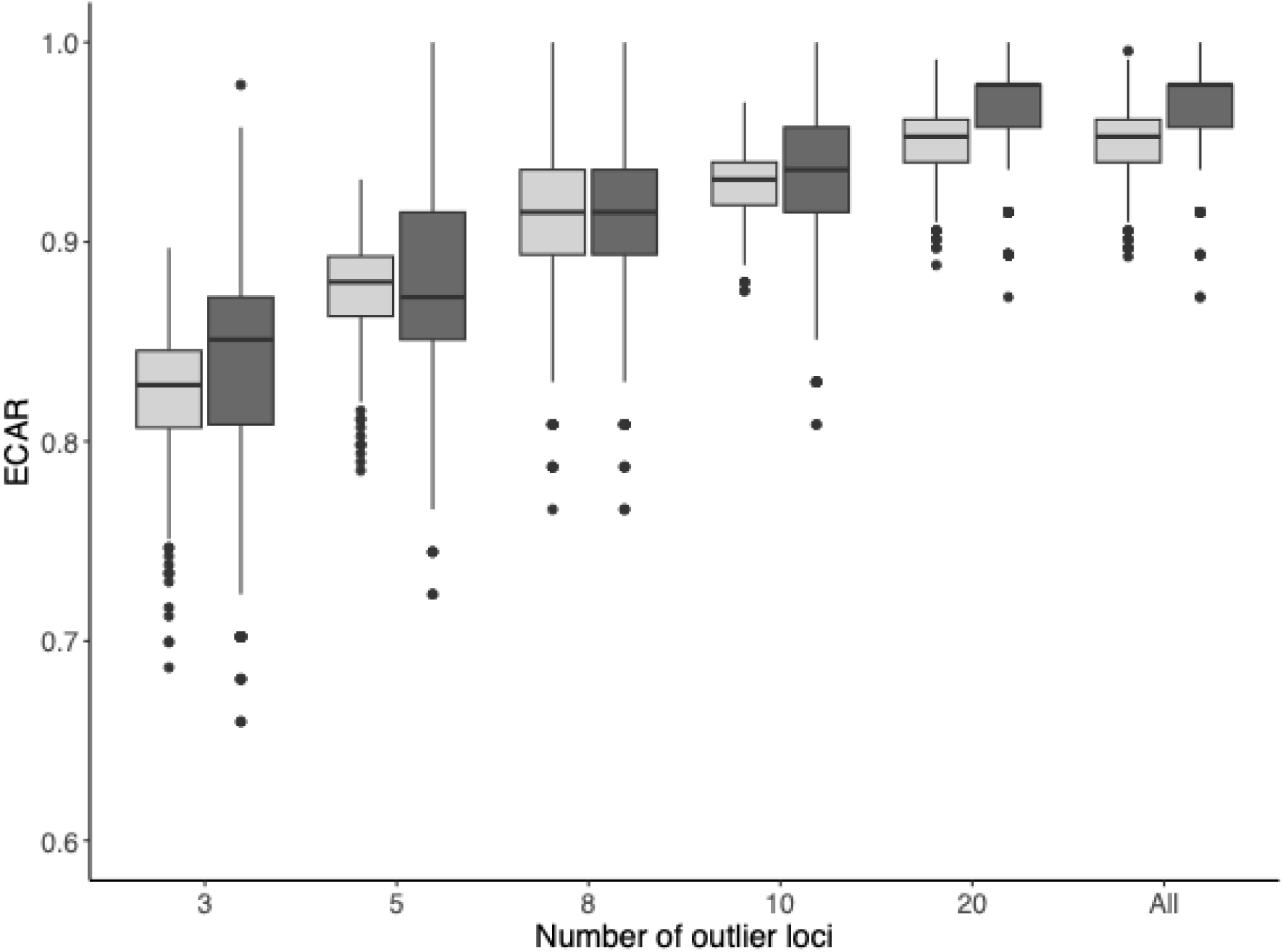
Estimated correct assignment rate (ECAR) for discriminating genetic groups via Monte Carlo cross-validation (1,000 iterations) with random sampling of training individuals at two levels (50%, light grey; 90%, dark grey) and across six levels of outlier loci (3, 5, 8, 10, 20, and all).

## Discussion

Population genomic analyses of Pacific cod across Japanese waters identified three distinct population groups: JBR, NHH, and WSJ (Fig. 4b). Demographic reconstruction using the PSMC approach revealed that the three genetic groups experienced distinct demographic histories since the LGP (Fig. 6).

Presently, the three groups are significantly differentiated (global *F*_ST_ = 0.056), with each exhibiting distinct genetic characteristics. The relatively high levels of differentiation between the WSJ group and other groups (*F*_ST_ = 0.061 for JBR vs. WSJ and 0.074 for NHH vs. WSJ) suggest genetic isolation of the WSJ group from the other groups, consistent with previous reports (Suda *et al*. 2017, Sakuma *et al*. 2019). In contrast, the lower differentiation level between the JBR and NHH groups (*F*_ST_ = 0.016) indicates a relatively high degree of genetic connectivity, possibly due to relatively recent divergence, historical migration, ongoing gene flow, and/or weak genetic drift. Nucleotide diversity (π) was highest in the JBR group, followed by the NHH and WSJ groups (Table 2; Supplementary Fig. S8). This pattern parallels the order of their recent historical *N*_e_ values (Fig. 6) and breadth of their geographic distribution ranges (Fig. 4b). The genome-wide patterns of Tajima’s D, an indicator of departure from the standard neutral demographic model, showed a consistent negative skew across all groups, with the strongest skew observed in the JBR group, followed by the NHH and WSJ groups (Table 2; Supplementary Fig. S9). These patterns are most plausibly explained by rapid population expansion, which increases the proportion of rare variants and elevates Watterson’s θ relative to π. Demographic history analysis further supported this interpretation, indicating varying degrees of recent expansion among groups (JBR > NHH > WSJ; Fig. 6). Taken together, our findings suggest that these three genetic groups follow distinct demographic trajectories.

The NHH group, related to the Mutsu Bay spawning aggregation known for strong natal homing (Fukuda *et al*. 1985, Miura *et al*. 2019), was identified for the first time in this study using genetic markers. NHH was exclusively detected in Mutsu Bay and adjacent waters along the northernmost Honshu on both the Sea of Japan and Pacific Coasts and also preferentially found along the Pacific Okhotsk Coast of Hokkaido (Fig. 4b). This spatial pattern broadly agrees with long-term tagged-recapture studies spanning over 30 years (Miura *et al*. 2019), which reported most recaptures in Mutsu Bay (77.9%), followed by the Pacific Coast of Hokkaido (14.3%) and Sea of Japan Coast of Hokkaido (4.3%). These findings suggest that the feeding grounds of the Mutsu Bay aggregation primarily encompass the Pacific Coast of Hokkaido, with some extension to the Sea of Japan Coast of southern Hokkaido. The recapture rate along the northern Honshu Coast was very low (1.3%), indicating limited feeding migration into this region. In contrast, genetic data showed that the NHH group was predominant in the northernmost part of Honshu on both the Sea of Japan and Pacific Coasts (Supplementary Fig. S13). These discrepancies may reflect spatiotemporal dynamics in population structure, as tagging–recapture surveys were conducted from 1979 to 2017, whereas genetic samples were collected from 2019 to 2022. Long-term genetic monitoring of the Atlantic cod *G. morhua* has revealed that the contributions of different stocks to mixed aggregations vary across years (Dahle *et al*. 2018). Therefore, spatiotemporal genetic monitoring is valuable for testing the hypothesis of fluctuations in Pacific cod population structure in Japanese waters. Caution should be exercised because some samples in this study (e.g., ID23 in Supplementary Fig. S13) were collected using minor fishing gear during a limited season, which may not fully represent the local population composition.

For future applications of GSI between the JBR and NHH groups, our resampling-based cross-validation suggests that over 500 neutral SNPs are necessary to achieve a discrimination accuracy > 0.9, whereas only over eight outlier SNPs are sufficient to achieve comparable accuracy (Fig. 7). Candidate outlier SNPs for practical implementation are listed in Supplementary Table S2. To evaluate the stability of these markers, we extended the resampling cross-validation procedure (Chen *et al*. 2017) by incorporating two different outlier detection methods: OutFLANK and pcadapt. This approach allowed us to assess whether the identified outliers consistently function as discriminative markers across resampling trials. With training data proportions of 50 and 90%, we detected 8 and 18 outlier SNPs, respectively, with a detection frequency ≥ 0.7 (i.e., each SNP was identified as an outlier in at least 70% of the 1,000 resampling trials). Functional annotation revealed that most outliers were found in introns or upstream regions, specifically with or near 5′-untranslated regions, of protein-coding genes and non-coding RNAs, including cytochrome P450 2J4-like (Supplementary Table S2). These results suggest that a relatively small set of carefully selected outlier SNPs associated with putative functional genes can effectively discriminate between the NHH and JBR groups, highlighting the practical utility of outlier markers for GSI.

IBD was observed among Pacific cod in Japanese waters (Fig. 5), consistent with previous reports from northeastern Pacific (Cunningham *et al*. 2009, Drinan *et al*. 2018). IBD was evident across Japanese waters and within the JBR, but not within the NHH. This suggests that gene flow is generally restricted by geographic distance across Japanese waters, but not within the NHH. This pattern implies that natal homing strongly influences genetic similarity within the NHH. However, individuals of the same genetic group do not necessarily spawn at the same location; effective gene flow between Mutsu Bay and other spawning grounds could maintain a shared genetic composition. The absence of IBD within the NHH does not exclude the possibility of additional spawning sites outside Mutsu Bay. Currently, Mutsu Bay is the only spawning site clearly documented within the NHH range. Historically, Kikonai Bay (on the Pacific Coast of Hokkaido, across the Tsugaru Strait from Mutsu Bay) has been recorded as a spawning site (Fujisawa and Natsume 1995); however, its current status remains unclear. This highlights the need to further investigate potential spawning grounds along the Hokkaido Coast.

Pacific cod is widely distributed across the North Pacific, with more than 10 putative stocks identified around Asia alone (Bakkala 1984). The observed IBD patterns across Japanese waters suggest effective gene flow at the regional scale, implying that gene flow also occurs between Japanese waters and adjacent regions. In this study, we incorporated two individuals from the Yellow Sea of China using publicly available data. Previous studies have shown that Pacific cod in the WSJ, roughly corresponding to the WSJ Coast of Honshu management unit, are genetically distinct from those in the northern Sea of Japan (Suda *et al*. 2017, Sakuma *et al*. 2019). Our results similarly detected genetic composition of the Chinese samples resembling that of the WSJ group, supporting a genetic connectivity between the WSJ and Yellow Sea. In contrast, the coastal waters around the Korean Peninsula appear to harbour more complex population structures (Kim *et al*. 2010, Gwak and Nakayama 2011, Fisher *et al*. 2022). Interestingly, the LD decay pattern in the WSJ group indicated potential population subdivision (Supplementary Fig. S8). Because these waters represent the southernmost limit of this species distribution, population structure may be more intricate, warranting further investigation. This study did not include samples from northern regions, such as Russian waters or other adjacent northern areas. Although PCA and DAPC are robust methods to explore genetic structure and do not rely on Hardy–Weinberg equilibrium assumptions, their ability to fully capture population differentiation remains limited when genetically distinct populations are not sampled. In such cases, the inferred structure may be oversimplified, potentially obscuring important patterns of diversity and connectivity. This underscores the critical importance of comprehensive and geographically representative sampling, particularly in peripheral regions, where unsampled populations may harbour unique genetic compositions that influence the overall structure. Population genomics research on Pacific cod has advanced considerably in the eastern North Pacific (Drinan *et al*. 2018, Sakuma *et al*. 2019, Spies *et al*. 2020, Spies *et al*. 2021, Spies *et al*. 2022b), and sequence data are becoming increasingly accessible through public databases. Future studies should evaluate stock structure from a broader perspective, encompassing the entire North Pacific distribution range. Such efforts are essential for a more comprehensive understanding of Pacific cod as a fishery resource and development of effective and sustainable management strategies.

## Acknowledgements

This study was conducted as part of the Marine Fisheries Stock Assessment and Evaluation of Japanese Waters program of the Fisheries Agency of Japan. We are grateful to all collaborators and coworkers for sample collection. We would like to thank Kunihiro Fujiwara, Osamu Sakai, and Kohei Hamabe for their valuable comments on an earlier version of the manuscript. The authors used Microsoft Copilot and OpenAI ChatGPT to check grammar and spelling and improve readability and language.

## Author contributions

ASH, KS, TA, and SNC designed the study. KS and SNC collected specimens and performed laboratory experiments. ASH and SNC performed bioinformatics analyses. ASH wrote the original draft of the manuscript, conducted all statistical analyses, and contributed to visualisation and data presentation. All authors reviewed and edited the manuscript and approved the final submission.

## Data availability

Raw FASTQ files of GRAS-Di and WGS data have been deposited in DRA/SRA/ERA (PRJDB18062 and PRJDB38066, respectively; see Supplementary Table S3 for details). All codes used for variant calling and resampling-based cross-validation procedures are available at https://github.com/akihirao/Gma_GRASDi.

