## Supplementary materials for "Genetic population structure and demographic history of Pacific cod in Japanese waters: Implications for stock identification using SNP markers"

### Supplementary Note

#### GRAS-Di sequencing

DNA was extracted from muscle tissues preserved in 6 M TNE–urea buffer (Asahida *et al.* 1996) using a Maxwell® RSC Blood DNA Kit (Promega Corporation, Madison, USA) following manufacturer's instructions. The GRAS-Di method—a reduced representation sequencing approach using a two-step PCR library with random primers (Enoki *et al.* 2018, Enoki *et al.* 2019) —was employed to identify genome-wide SNPs. Library construction and sequencing were outsourced to GeneBay Inc. (Yokohama, Japan) with minor modifications. Due to specific amplification observed in preliminary experiments, the NE10_388 primer was removed from the 1st-PCR primer set. The first library, which included 435 individuals, and the second library, which included 80 individuals, were independently sequenced on an Illumina NovaSeq 6000 platform (150 bp, paired-end).

#### Variant calling procedure for GRAS-Di dataset

All custom scripts and commands for variant calling and SNP genotyping are available at https://github.com/akihirao/Gma_GRASDi. The variant calling procedure was adapted from the germline short-variant discovery workflow in the GATK Best Practices recommendations (DePristo *et al.* 2011). Raw reads from GRAS-Di samples were cleaned by removing adapter sequences and low-quality sequences using NGmerge v.0.3 (Gaspar, 2018), Trimmomatic v0.36 (Bolger et al, 2014), and fastp v.0.22.0 (Chen et al, 2018). The cleaned reads were aligned to the *Gadus macrocephalus* reference genome (GenBank accession: GCF_031168955.1) using BWA-mem2 v2.2.1 (Vasimuddin *et al.* 2019), and multi-mapped reads were removed using SAMtools v1.16.1 (Danecek *et al.* 2021). Both variant and non-variant sites were called by using the "---include-non-variant-sites" option in the GATK GenotypeGVCFs command, by followed by the removal of indels, tri- or tetra-allelic SNP, low-confidence sites (QD < 2.0, MQ < 40.0, MQRankSum < -12.5, ReadPosRankUsm < -8.0, SOR > 4.0, and ExcessHet > 13.0), and low-confidence genotypes (DP < 20, GQ or RGQ < 30). The remained high-quality bi-allelic SNPs and non-variants were subjected to multi-step filtering to retain loci with a missing rate < 0.1 using PLINK v1.90 (Purcell et al. 2007) (see Hirao et al. 2024 for details). Bi-allelic SNPs (including singletons) and non-variants with a missing rate < 0.1 were used for estimating nucleotide diversity and analyzing LD decay (Dataset 1). For population structure analyses, SNPs with a minor allele frequency (MAF) > 0.05 were retained after linkage disequilibrium (LD) pruning ($r^2 < 0.1$) using PLINK (Dataset 2). To check for potential cross-contamination and unintended sample duplicates, pairwise kinship coefficients among individuals were calculated using KING-robust methods using PLINK.

### Procedure for generating Dataset 3

Dataset 3 involved integrating two resequenced samples from the Yellow Sea, China (DRA/SRA/ERA accession no.: SRR21531029, SRR17394978) with the GRAS-Di dataset (Dataset 2) collected from Japanese coastal waters. Raw reads from the two sequenced samples were cleaned by removing adapter sequences and low-quality sequences using fastp v.0.22.0 (Chen *et al.* 2018). The cleaned reads were then aligned to the *Gadus macrocephalus* reference genome (GenBank accession: GCF_031168955.1) using BWA-mem2 v2.2.1 (Vasimuddin *et al.* 2019), and multi-mapped reads were removed using SAMtools v1.16.1 (Danecek *et al.* 2021). Variant calling followed the germline short variant discovery workflow in GATK ver. 4.5.0.0 (McKenna *et al.* 2010). Both variant and non-variant sites were called by using the "---include-non-variant-sites" option in the GATK GenotypeGVCFs command, followed by the removal of indels, tri- or tetra-allelic SNP, low-confidence sites (QD < 2.0, FS > 60.0, MQ < 40.0, MQRankSum < -12.5, ReadPosRankUsm < -8.0, SOR > 4.0, and ExcessHet > 13.0), and low-confidence genotypes (DP < 20, GQ or RGQ < 30). The high-quality bi-allelic SNPs and non-variants from the two resequencing samples were cross-referred and merged with those from the GRAS-Di samples (Dataset 2) using vcf-merge function in VCFtools ver0.1.16 (Danecek *et al.* 2011). SNPs with a minor allele frequency (MAF) > 0.05 and missing data rate < 0.1 were filtered prior to LD pruning ($r^2 < 0.1$) using PLINK, resulting in Dataset 3 for population analyses.

### Phylogenetic analysis

Phylogenetic relationships among individuals sampled from Japanese coastal wataers and the Yellow Sea (Dataset 3) were inferred using the maximum-likelihood method implemented in IQ-TREE v2.2.2.7 (Minh *et al.* 2020), applying the GTR+ASC model with 1,000 ultrafast bootstrap replicates. The resulting tree was rooted with an individual from Kodiak Island, Alaska, which served as the reference genome assembly (BioSample ID: SAMN3376189). The all-site genotype of the Alaska individual was determined by aligning the individual's alternative haplotype assembly (ASM3119387v1: https://www.ncbi.nlm.nih.gov/datasets/genome/GCA_031193875.1) to its reference assembly (ASM3116895v1: https://www.ncbi.nlm.nih.gov/datasets/genome/GCF_031168955.1) using minimap2 (Li 2018). The genotype of the Alaska individual was then cross-referenced and merged with those from Dataset 3 using vcf-merge function in VCFtools ver0.1.16 (Danecek *et al.* 2011) for phylogenetic analylsis.

### Whole-genome sequencing and PSMC analyses

Whole-genome resequencing was performed for one representative individual from each of the three groups identified (See results in main text). Two individuals were sequenced using the Illumina NovaSeq 6000 platform (150 bp, paired-end), and one individual using the DNBSEQ-T7 platform (150 bp, paired-end). Read filtering, alignment, and variant calling followed the procedure described in the previous section. Aligned BAM files, from which multi-mapping reads and repeat element regions had been filtered out, were used for pairwise sequentially Markovian coalescent (PSMC) analyses (Li and Durbin 2011). Repeat elements were predicted *de novo* using RepeatModeler v2.0.4 and masked against the reference genome using RepeatMasker v4.1.4 (https://www.repeatmasker.org/).

PSMC input files were generated using BCFtools mpileup from the aligned BAM files, applying a minimum mapping quality threshold of 30. Sites with read depths greater than twice or less than one-third of the average depth were excluded. PSMC was run for 30 iterations, with a maximum coalescent time ($t$) set to 15, and the mutation-to-recombination rate ratio ($r$) of 5. Atomic time intervals ($p$) were specified as '6 + 23×2 + 6'. These settings followed Li and Durbin's guideline that each time interval contain more than 10 inferred recombination events (https://github.com/lh3/psmc). As preliminary tests, we conducted PSMC runs using an alternative setting for $p$ ('4+ 10×1 + 20×2 + 4 + 6') with the other parameters retained—previously applied for comparative dehomographic histories across 22 teleost species (Li *et al.* 2021)— to confirm that estimated population dynamics were not depend on specific settings for $p$. Furthermore, we preliminary tested another alternative setting of $p$ ('2 + 2 + 2 + 23×2 + 6'), in which the first time interval was split, to confirm the specifit artifact pattern—recent population peaks followed by population collapse (Hilgers *et al.* 2025)—did not occur. Variance in PSMC estimates was assessed using 100 bootstrap replicates by resampling 500-kb genome segments. Effective population size ($N_e$) and time in years were scaled using a mutation rate of $1.64 \times 10^{-8}$ per site per generation (estimated for Atlantic cod; Matschiner *et al.* 2022) and a generation time of 9 years (see next section). To assess sensitivity to generation time, minimum and maximum values (4 and 14 years, respectively; see next section) were also applied.

#### Generation time for Pacific cod

The species-specific generation time for Pacific cod was calculated using the equation of

Pacoureau *et al.* (2021):

$$GT = (A_{max} + A_{mat})\, z + A_{mat}$$

where the age at maturity ($A_{mat}$) was 3.6 years and the maximum age ($A_{max}$) was 14.3 years, both obtained from FishBase (Thorson *et al.* 2017), and the mortality-related coefficient (z) was conservatively set to 0.5 (Pacoureau *et al.* 2021). The estimated generation time was 8.7 years, which we rounded to 9 years for subsequent analyses. To assess the sensitivity of this assumed generation time of 9 years, minimum and maximum generation times of 4 and 14 years, respectively, were also evaluated. The minimum generation time was inferred based on the age at which 100% maturity is reached (Sakuma *et al.* 2025), while the maximum generation time was calculated using the maximum reported age of 25 years (Munk 2001), with the other parameters retained.

Asahida T, Kobayashi T, Saitoh K, *et al.* Tissue preservation and total DNA extraction form fish stored at ambient temperature using buffers containing high concentration of urea. *Fisheries science* 1996; 62: 727–730.

Chen S, Zhou Y, Chen Y, *et al.* fastp: an ultra-fast all-in-one FASTQ preprocessor. *Bioinformatics* 2018; 34: i884–i890. https://doi.org/10.1093/bioinformatics/bty560

Danecek P, Auton A, Abecasis G, *et al.* The variant call format and VCFtools. *Bioinformatics* 2011; 27: 2156–2158. https://doi.org/doi:10.1093/bioinformatics/btr330

Danecek P, Bonfield JK, Liddle J, *et al.* Twelve years of SAMtools and BCFtools. *Gigascience* 2021; 10: giab008. https://doi.org/doi:10.1093/gigascience/giab008

DePristo MA, Banks E, Poplin R, *et al.* A framework for variation discovery and genotyping using next-generation DNA sequencing data. *Nat Genet* 2011; 43: 491–498.

Enoki H, Takeuchi Y, Suzuki K. New genotyping technology, GRAS-Di, using next generation sequencer. In: *Proceedings of the Plant and Animal Genome Conferene XXVI, Sandiego, CA*, 2018, https://pag.confex.com/pag/xxvi/meetingapp.cgi/Paper/29067

Enoki H, Takeuchi Y, Suzuki K. Genotyping By Random Amplicon Sequencing-Direct, GRAS-Di. In: *Plant and Animal Genome XXVII Conference (January 12-16, 2019)*: PAG, 2019, https://pag.confex.com/pag/xxvii/meetingapp.cgi/Paper/33956

Hilgers L, Liu S, Jensen A, *et al.* Avoidable false PSMC population size peaks occur across numerous studies. *Current Biology* 2025; 35: 927–930. e923.

Li H, Durbin R. Inference of human population history from individual whole-genome sequences. *Nature* 2011; 475: 493–496. https://doi.org/10.1038/nature10231

Li H. Minimap2: pairwise alignment for nucleotide sequences. *Bioinformatics* 2018; 34: 3094–3100.

Li J, Bian C, Yi Y*, et al.* Temporal dynamics of teleost populations during the Pleistocene: a report from publicly available genome data. *BMC genomics* 2021; 22: 1–11.

Matschiner M, Barth JMI, Tørresen OK*, et al.* Supergene origin and maintenance in Atlantic cod. *Nature Ecology & Evolution* 2022; 6: 469–481.

McKenna A, Hanna M, Banks E*, et al.* The Genome Analysis Toolkit: a MapReduce framework for analyzing next-generation DNA sequencing data. *Genome Res* 2010; 20: 1297–1303. https://doi.org/10.1101/gr.107524.110

Minh BQ, Schmidt HA, Chernomor O*, et al.* IQ-TREE 2: New Models and Efficient Methods for Phylogenetic Inference in the Genomic Era. *Mol Biol Evol* 2020; 37: 1530–1534. https://doi.org/10.1093/molbev/msaa015

Munk KM. Maximum ages of groundfishes in waters off Alaska and British Columbia and considerations of age determination. *Alaska Fishery Research Bulletin* 2001; 8: 12–21.

Pacoureau N, Rigby CL, Kyne PM*, et al.* Half a century of global decline in oceanic sharks and rays. *Nature* 2021; 589: 567–571. https://doi.org/10.1038/s41586-020-03173-9

Sakuma K, Naito T, Kinoshita S*, et al.* Stock assessment and evaluation for Pacific cod of Sea of Japan Honshu stock (fiscal year 2025) (in Japanese). Marine fisheries stock assessment and evaluation for Japanese waters. Japan Fisheries Agency and Japan Fisheries Research and Education Agency, Tokyo. p. 1–31. FRA-SA2025-SC03-04. 2025.

Thorson JT, Munch SB, Cope JM*, et al.* Predicting life history parameters for all fishes worldwide. *Ecological Applications* 2017; 27: 2262–2276. https://doi.org/10.1002/eap.1606

Vasimuddin M, Misra S, Li H*, et al.* Efficient architecture-aware acceleration of BWA-MEM for multicore systems. In: *2019 IEEE International Parallel and Distributed Processing Symposium (IPDPS)*: IEEE, 2019, 314–324

Waples RS. Practical application of the linkage disequilibrium method for estimating contemporary effective population size: A review. *Mol Ecol Resour* 2024; 24: e13879. https://doi.org/10.1111/1755-0998.13879

### Supplementary Figures

**Figure S1**. Principal component analysis (PCA) of 496 Pacific cod individuals, showing PC1 vs. PC2 (top) and PC1 vs. PC3 (bottom). Colors and symbols indicate management units.

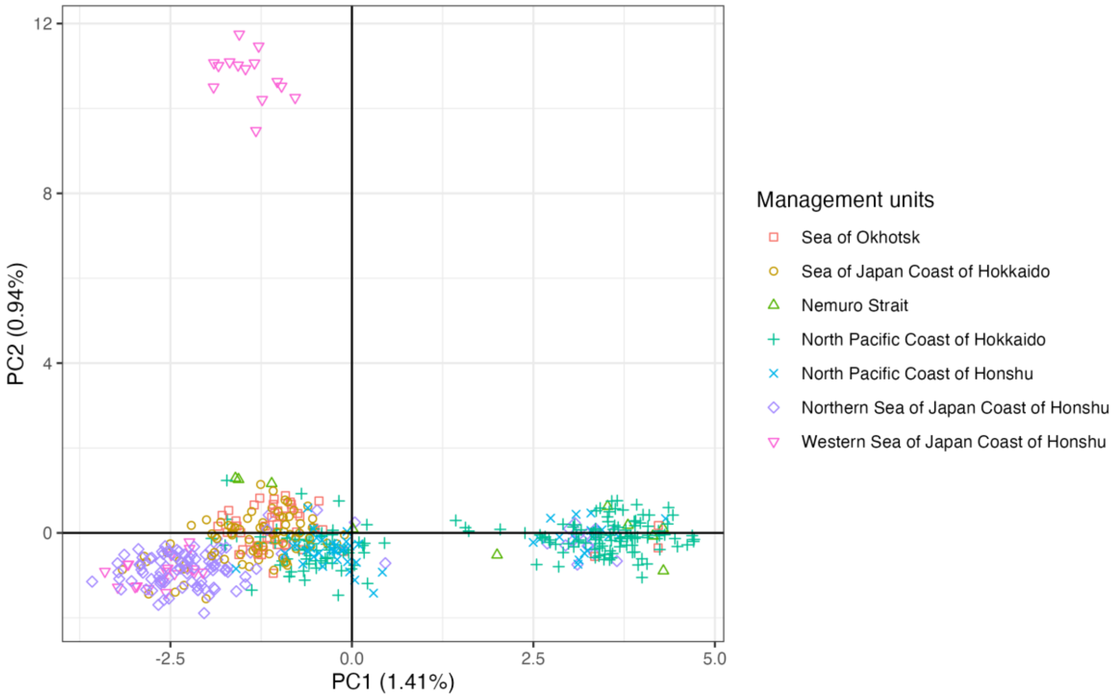


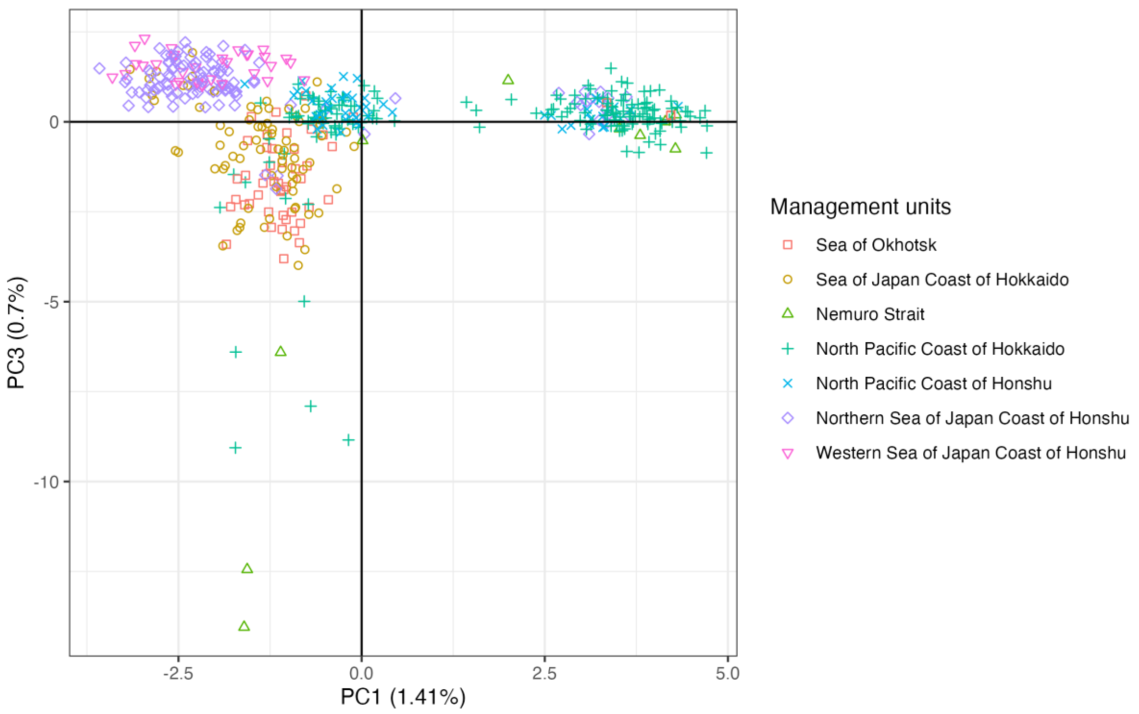

**Figure S2**. Screeplot from the Principal Component Analysis (PCA) displaying the number of principal components against their corresponding eigen values.

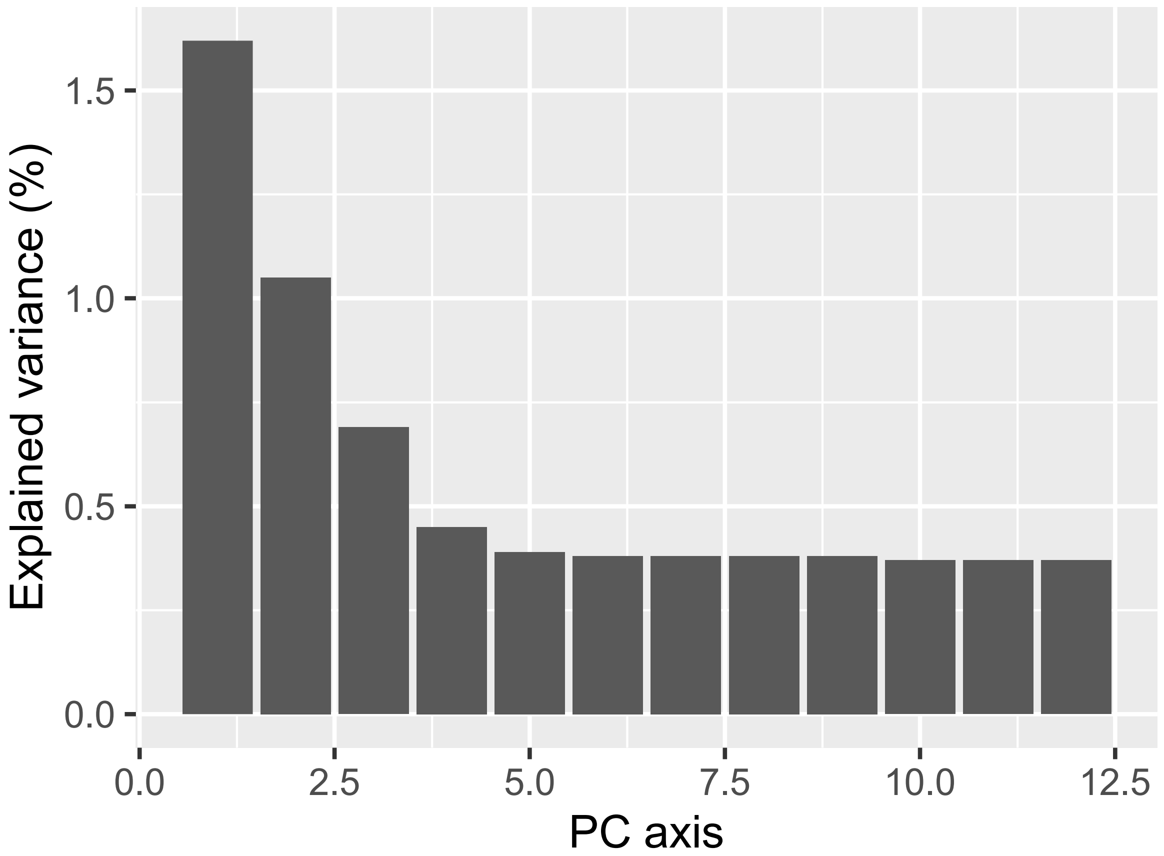

**Figure S3.** Principal component analysis (PCA) of 498 Pacific cod individuals, including 496 from Japanese coastal waters and 2 from the Yellow Sea, showing PC1 vs. PC2 (top) and PC1 vs. PC3 (bottom). Colors and symbols indicate management units.

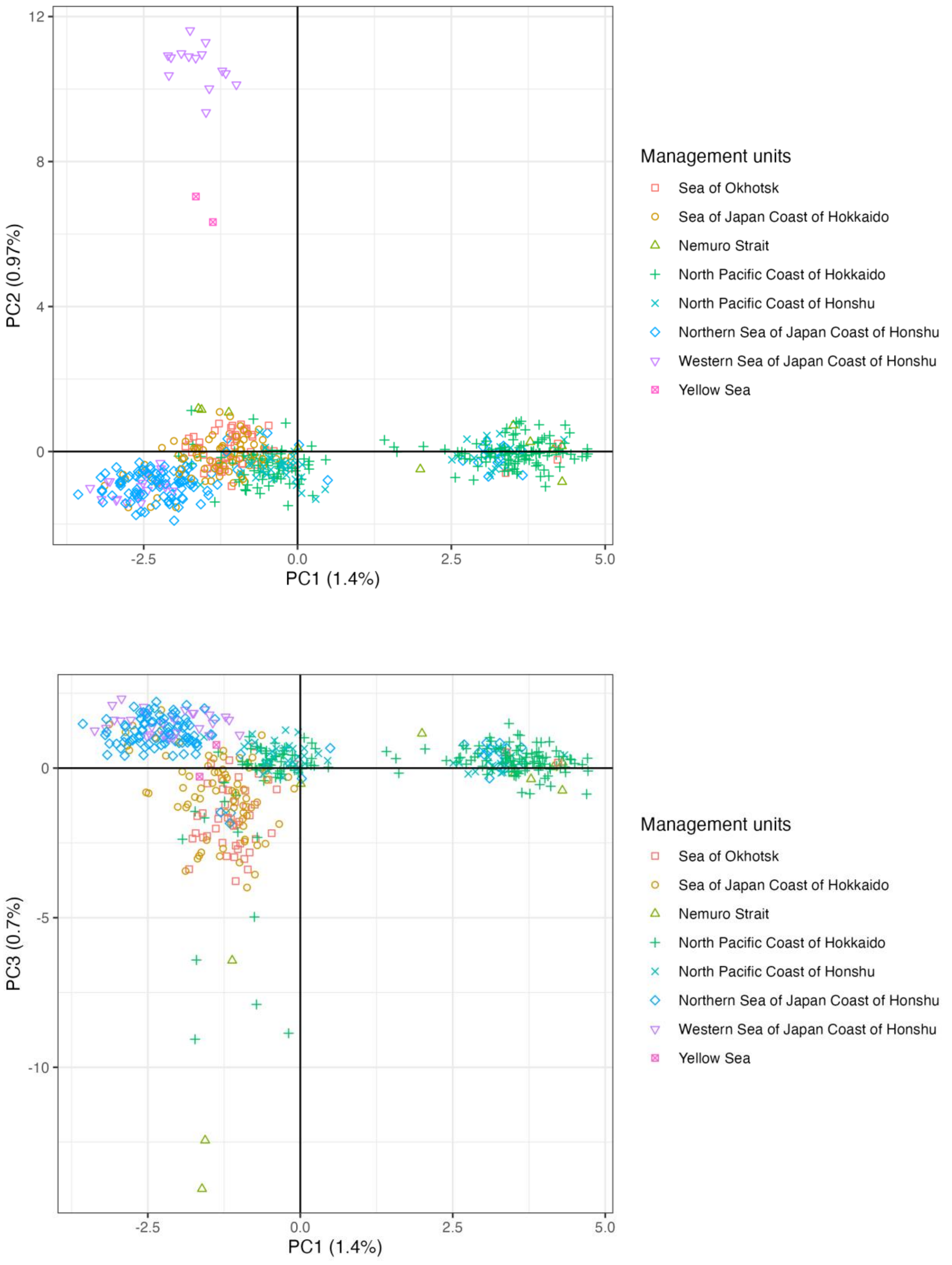

**Figure S4.** Discriminant Analysis of Principal Component (DAPC) for 498 Pacific cod individuals, including 496 from Japanese coastal seas and 2 from the Yellow Sea. Colors indicate groups identified by DAPC: yellow – Japanese Broad Range group (JBR); green – Northernmost Honshu–Hokkaido group (NHH); blue – Western Sea of Japan group (WSJ).

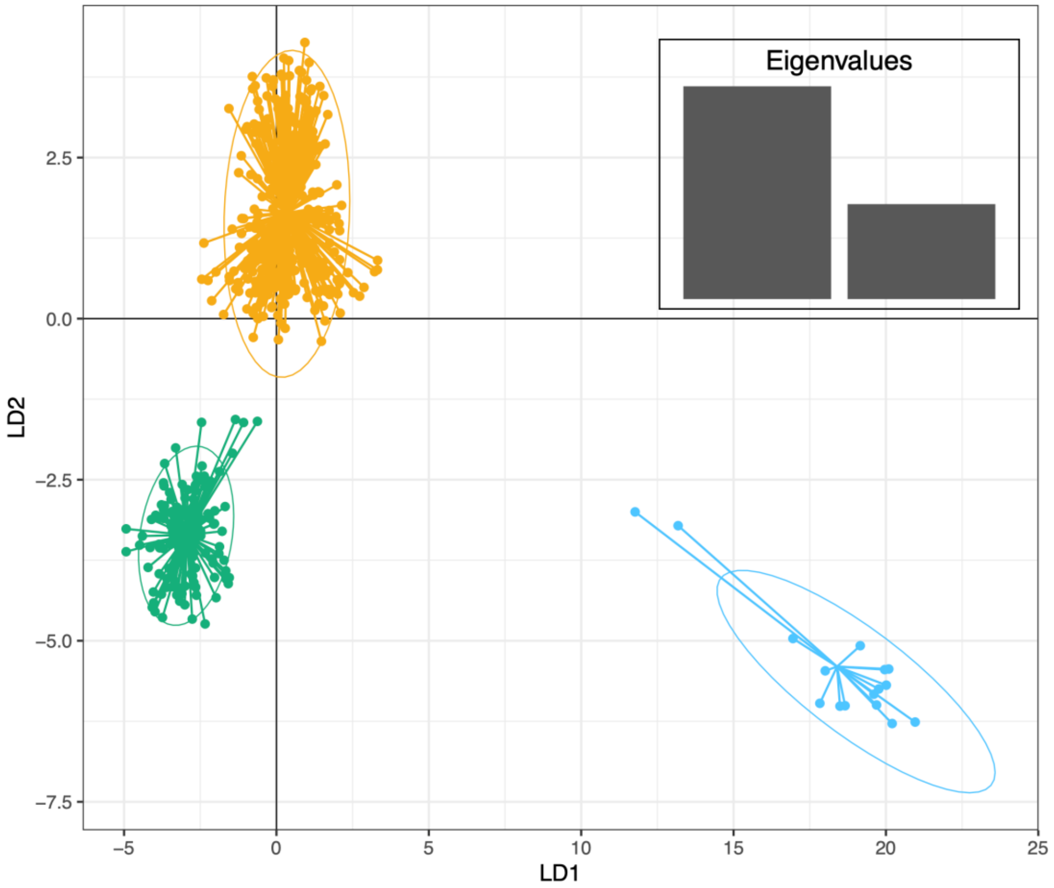

**Figure S5.** Map of Japanese coastal waters showing pie charts representing membership proportions to three genetic groups identified by discriminant analysis of principal components (DAPC) for 468 Pacific cod individuals, including two from the Yellow Sea. Colors indicate genetic groups: orange – Japanese Broad Range group (JBR); green – Northernmost Honshu–Hokkaido group (NHH); blue – Western Sea of Japan group (WSJ).

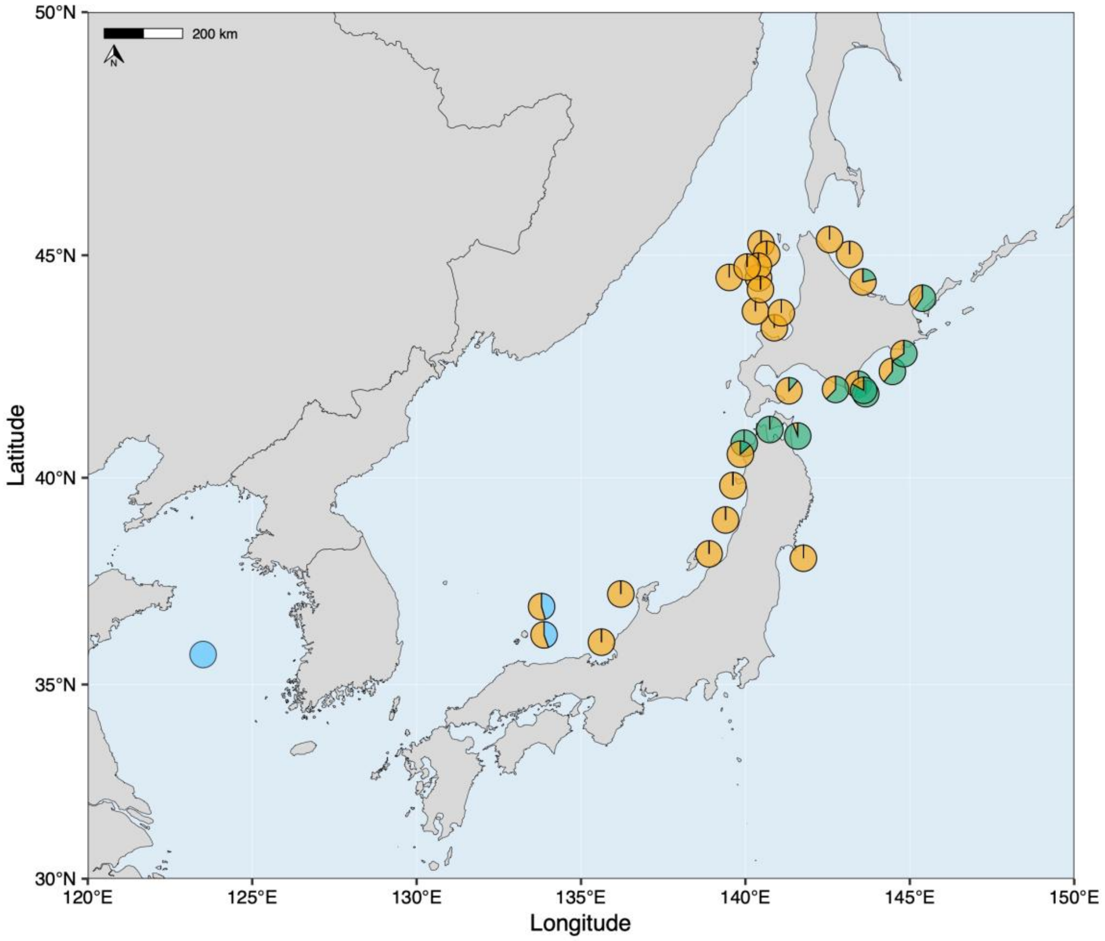

**Figure S6.** Maximum-likelihood phylogenetic tree of Pacific cod from Japanese coastal waters and the Yellow Sea, rooted with an individual from Kodiak Island, Alaska, which served as the reference genome assembly (BioSample ID: SAMN3376189). Colors and shapes indicate genetic groups: yellow circle – Japanese Broad Range group (JBR); green triangle – Northernmost Honshu–Hokkaido group (NHH); blue square – Western Sea of Japan group (WSJ); blue cross – Yellow Sea (CHN); grey circle – undetermined group (UND: posterior assignment probability < 0.95); open circle – Alaska. Nodes within the HPM and WSJ groups were each supported by 100% bootstrap replicates.

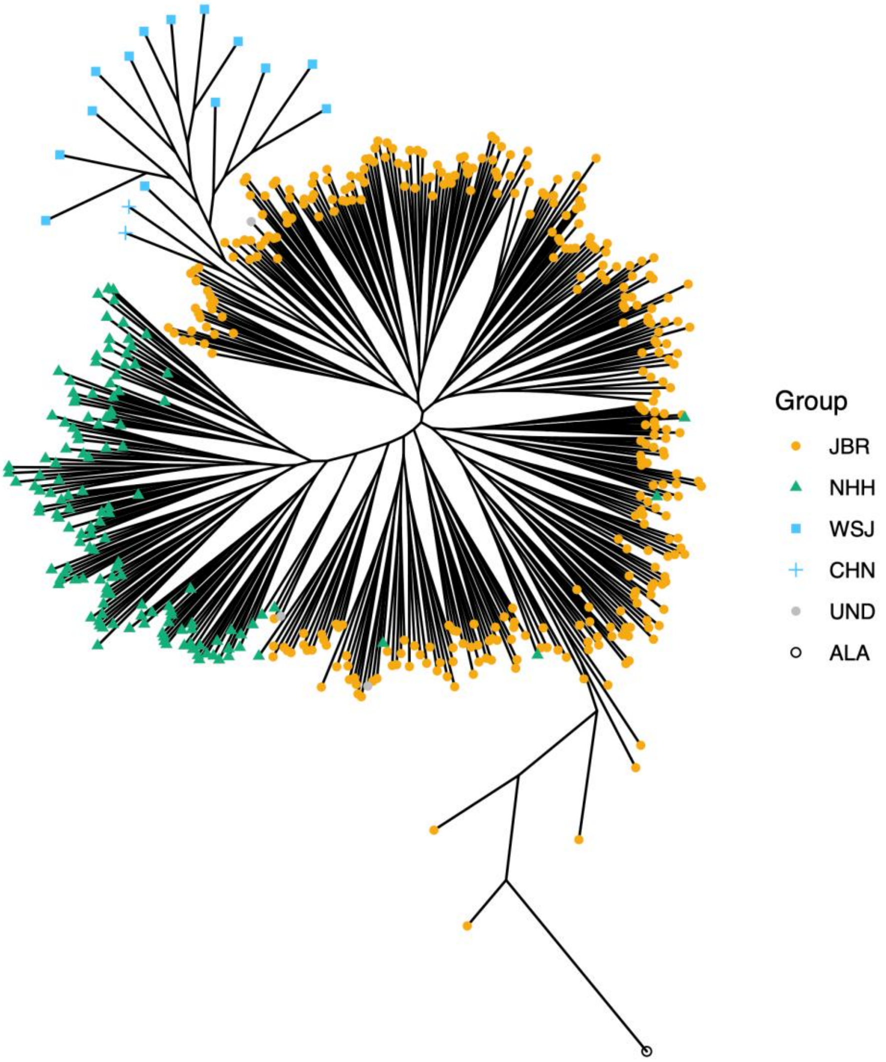

**Figure S7.** Nucleotide diversity ($\pi$) within three genetic groups identified by DAPC: Japanese Broad Range group (JBR); Northernmost Honshu–Hokkaido group (NHH); Western Sea of Japan group (WSJ). The levels of $\pi$ were significantly different between JBR and WSJ ($P < 0.05$ by window-blocked bootstrap test with Bonferroni correction), whereas no significant differences were observed among the other pairwise group comparisons.

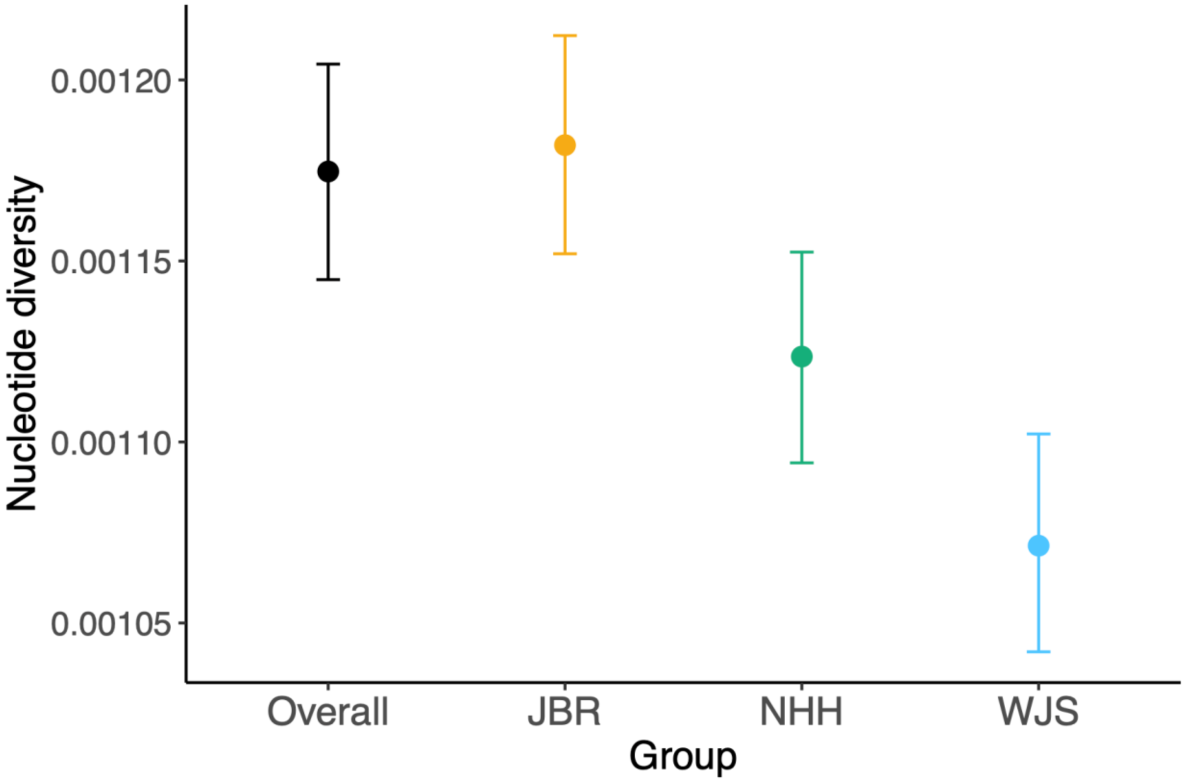

**Figure S8.** Linkage disequilibrium (LD) decay represented by average genotypic association coefficient ($r^{2\prime}$)—adjusted $r^2$ for finite sample size (Waples 2024)—as a function of inter-SNP distance.

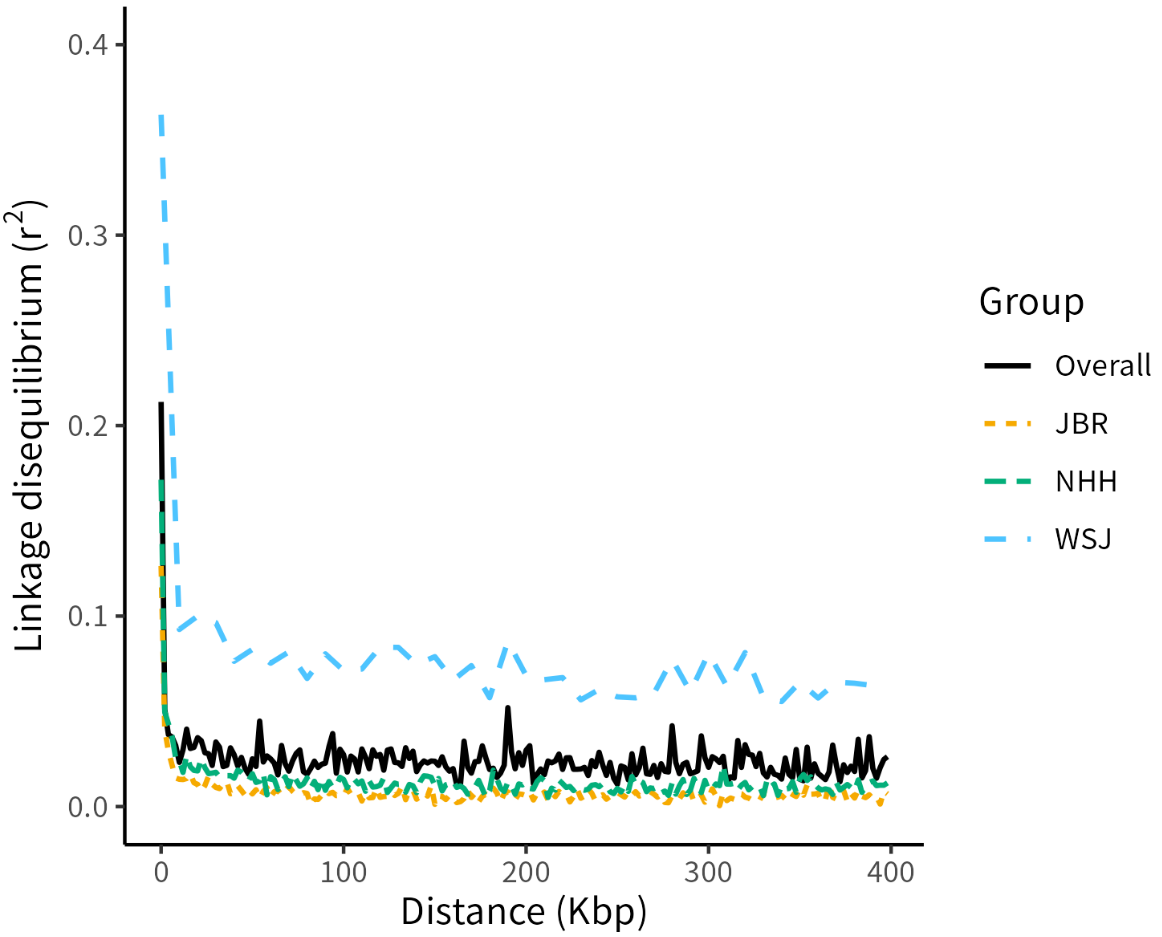

**Figure S9.** Genome-wide Tajima's D for three genetic groups identified by DAPC: Japanese Broad Range group (JBR); Northernmost Honshu–Hokkaido group (NHH); Western Sea of Japan group (WSJ). The distribution pattern of Tajima's D was negatively biased and differed significantt among groups ($P < 0.001$ for all pairs, using a linear mixed model with Student-t errors [glmmTMB], accounting for windows-block structure with Holm adjustment).

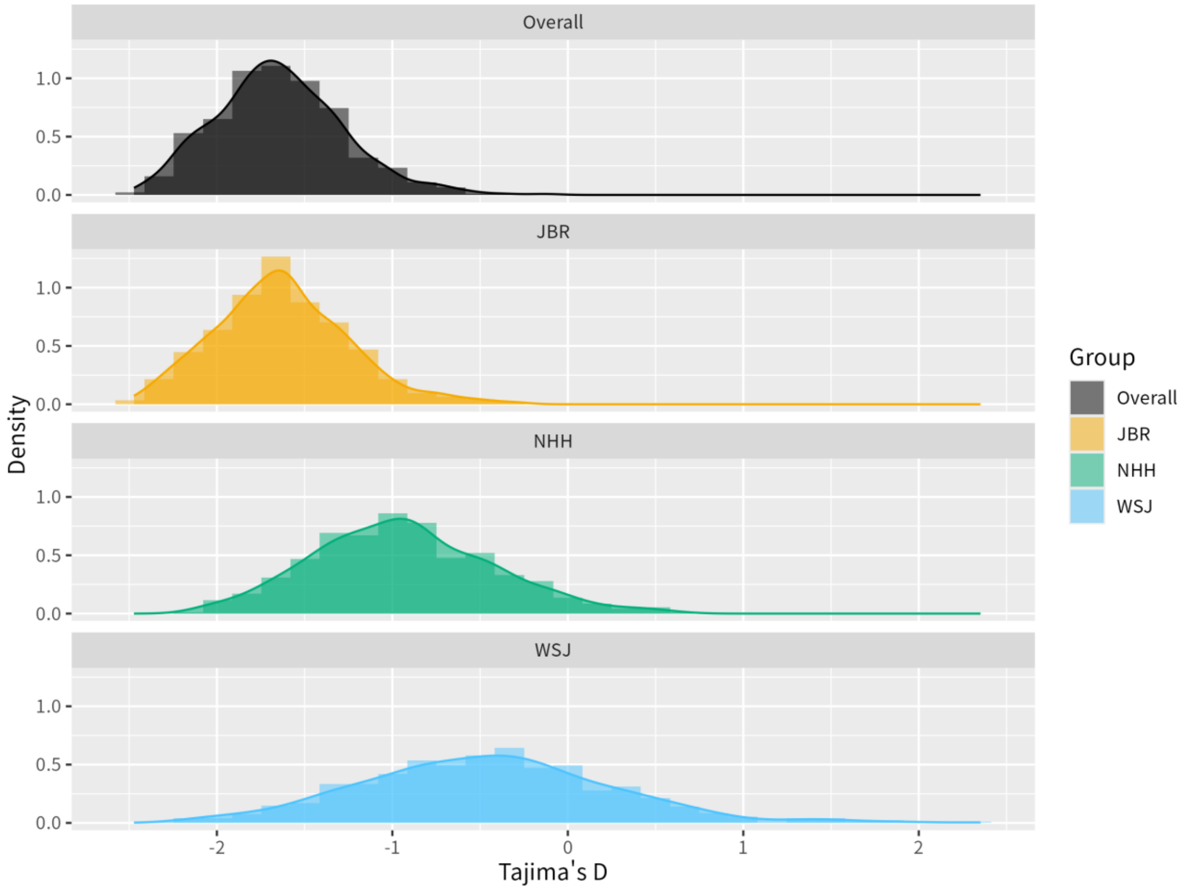

**Figure S10.** PSMC estimates based on whole-genome resequencing (WGS) data from one representative individual for each genetic group: Japanese Broad Range group (JBR: yellow), Northernmost Honshu–Hokkaido group (NHH: green), and Western Sea of Japan group (WSJ: blue). As part of sensitivity analyses, minimum and maximum assumed generation times of 4 and 14 years were used to scale the time axis in the upper and lower panels, respectively. Bold lines show estimates from observed data, and thin lines represent 100 bootstrap replicates. The shaded area indicates the Last Glacial Period (LGP).

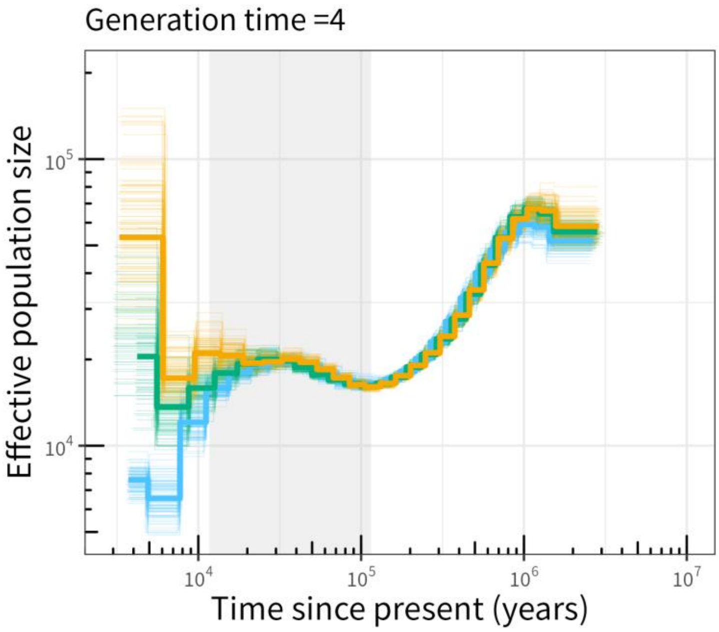


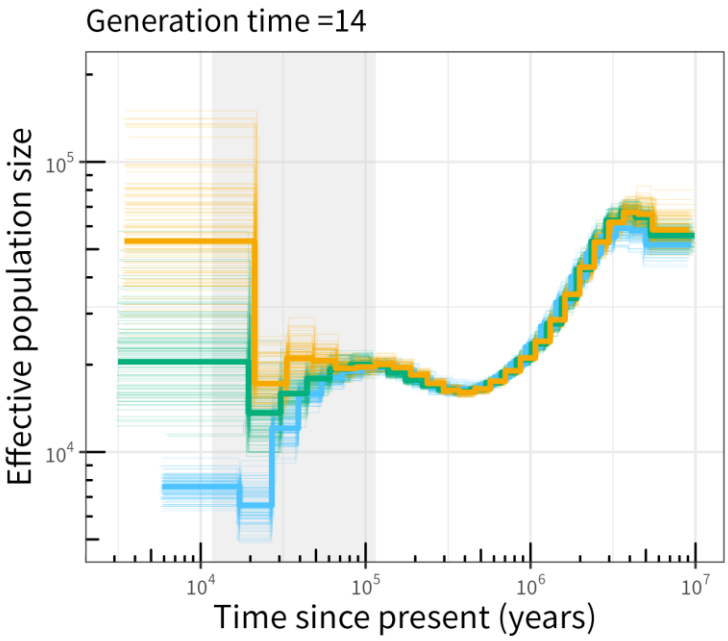

1. **Figure S11.** Estimated Correct Assignment Rate (ECAR) for discriminating genetic groups via Monte–Carlo cross-validation (1,000 iterations) with random sampling of training individuals at two levels (50%, light grey; 90%, dark grey) and across six levels of background SNPs (10, 100, 300, 500, 1000, and all).

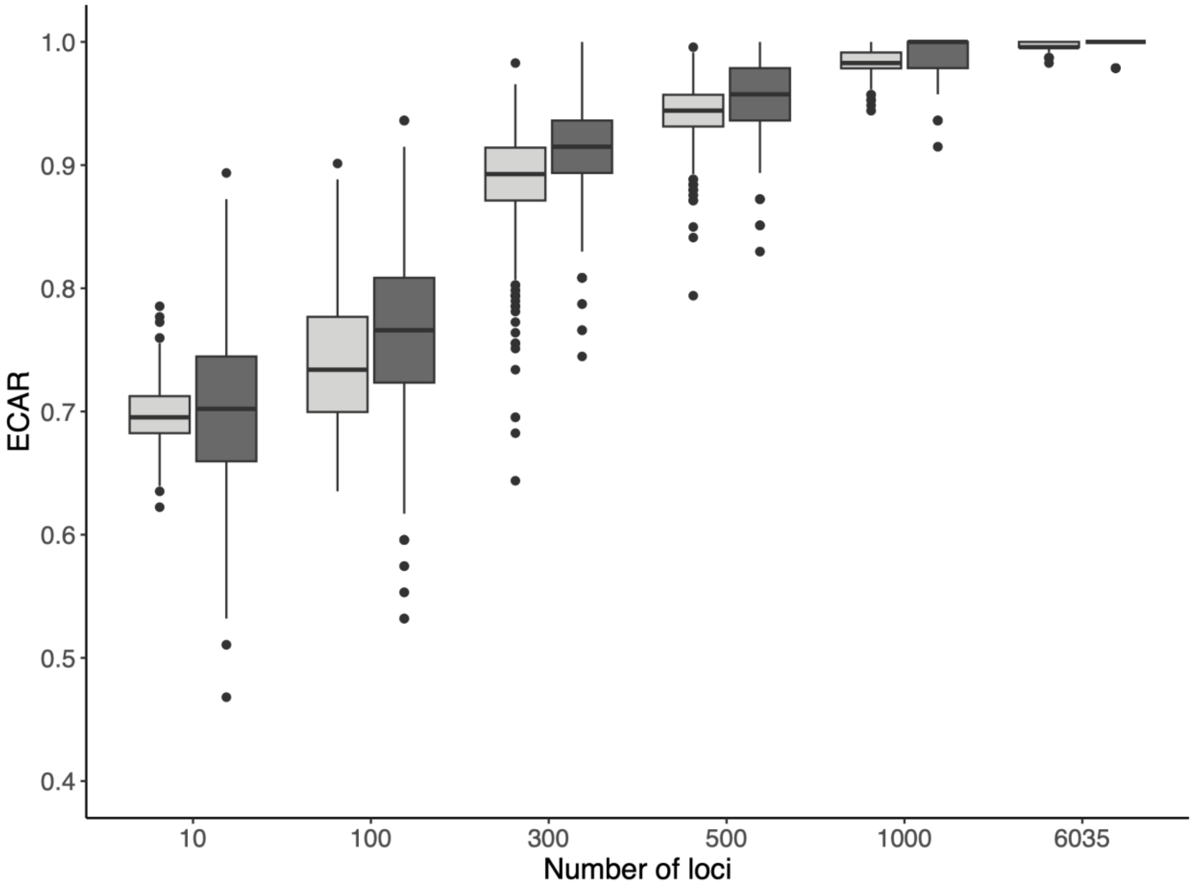

**Figure S12.** Frequency of 151 outlier SNPs obtained from all cross-validation procedures (1,000 iterations), with random sampling of training individuals at two levels (50% and 90%) and six thresholds of outlier loci (top 3, top 5, top 8, top 10, top 20, and all). Outlier SNPs are arranged along chromosomal positions on the x-axis.

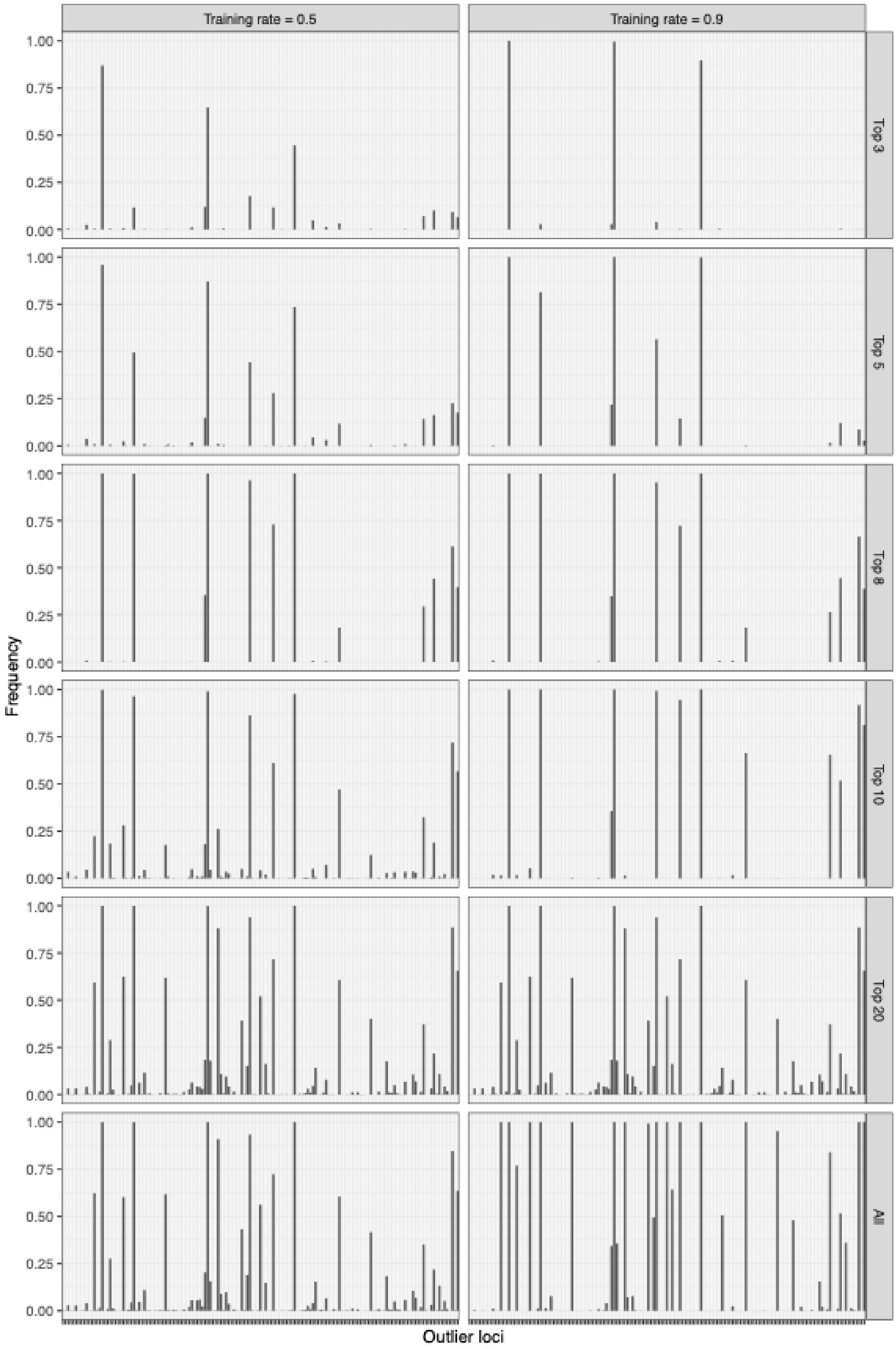

**Figure S13.** Distribution map of three genetic groups identified by discriminant analysis of principal components (DAPC). Colors indicate genetic groups: yellow – Japanese Broad Range group (JBR); green – Northernmost Honshu–Hokkaido group (NHH); blue – Western Sea of Japan group (WSJ).

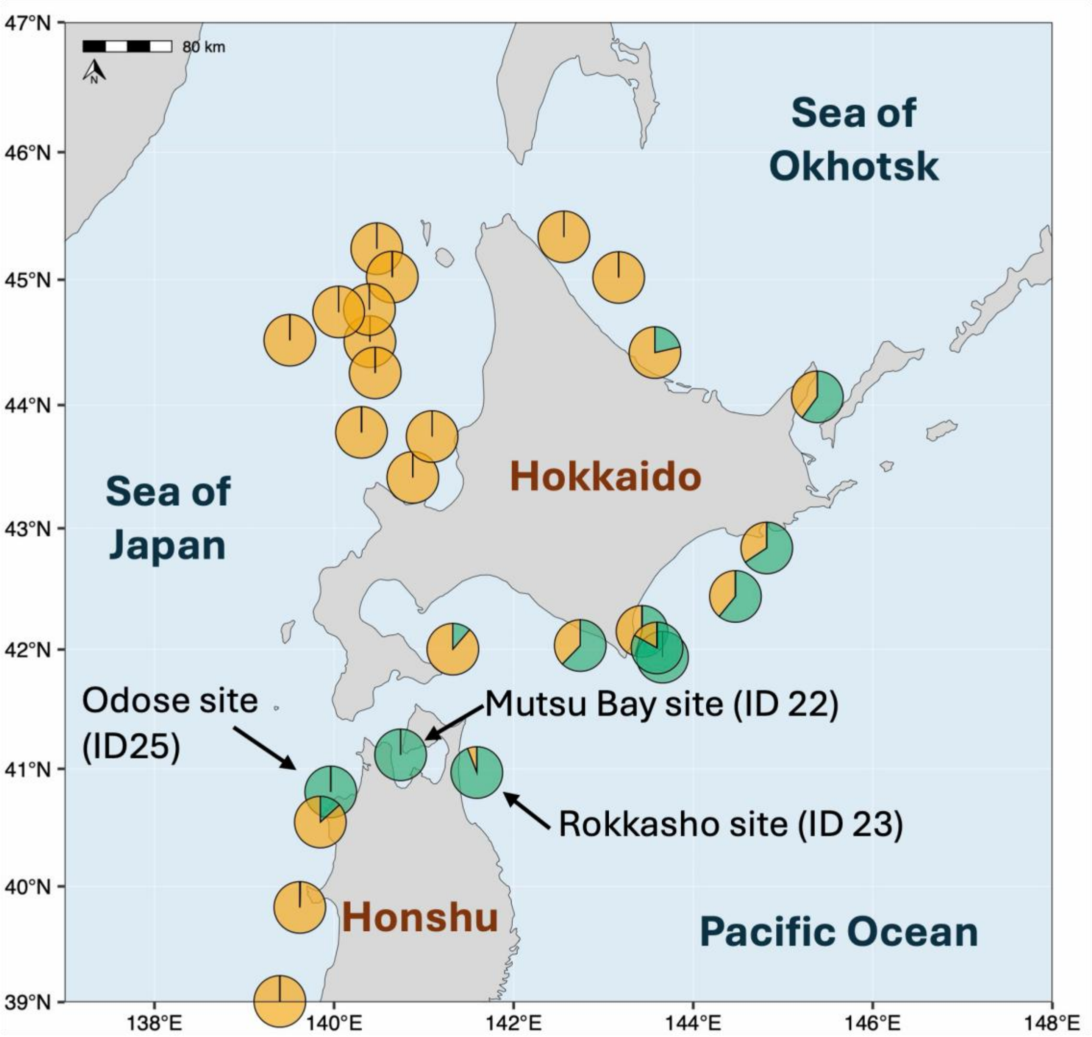
